# Common marmosets use body posture as multi-functional signal to solicit, maintain, and modify social play

**DOI:** 10.1101/2024.08.14.607991

**Authors:** Jessie E.C. Adriaense, Erik J. Ringen, Atsushi Ohashi, Judith M. Burkart

## Abstract

Social play is a highly active social interaction, characterized by rapid exchanges of various behaviors with multiple partners. Many primates use bodily expressions during social play, yet the potential signaling function of these expressions remains unclear. This study investigated whether common marmosets (*Callithrix jacchus)* use body posture as signal to regulate play. We recorded play within three captive common marmoset family groups using multiple cameras simultaneously to capture the fast-paced and high frequency behaviors. Three distinct signals (i.e. supine, hide, stalk) and six distinct play types (i.e. wrestle, chase, pounce, touch, catch, pull) were identified. We used a multi-state time-to-event model to analyze the sequences of play, including short-and long-term transitions between different states (i.e. signal, play, or rest/nothing). Our data-driven approach accounted for uncertainty in the duration of play bouts, using probabilistic classification rather than arbitrary bout thresholds. The resulting classifications allowed us to assess the social function of signals by comparing play behavior to a resting state baseline. We found that the presence of a signal: (1) increases the probability to play; (2) extends the duration of play; (3) leads to more diverse play; and (4) increases the probability of play fighting. Marmosets also show turn-taking of signaling and initiating subsequential play. These results show that marmosets use postures as communicative signals to initiate and change play dynamics, and thereby establish a mutual understanding of the joint action. The two-fold contribution of this study concerns novel analytical methods and a deeper conceptual understanding of primate communication. Play and its signals are important elements in the evolution of language, and our research contributes to its further understanding.

## INTRODUCTION

Animal play concerns multiple behaviors which vary greatly within and between species (Chalmers & Locke-Haydon, 1981; Held & Špinka, 2011). Play seems to be prominent in many species (Graham & Burghardt, 2010) with most occurrences observed in mammals and birds (Osvath & Sima, 2014; Kaplan, 2024), though a species bias in play research might reveal an underrepresentation. In particular, primates show the most play of all mammalian species (Burghardt, 2005) with the majority of play performed by juveniles and less common occurrences in adults (Ciani *et al*., 2012; Asensio *et al*., 2022). Overall, play in animals can be categorized as solitary, such as object or locomotor play, or social, which is directed toward another individual (Fagen, 1982; Held & Špinka, 2011). Research on social play has largely focused on its adaptive value, yet, despite many theories, the precise costs and benefits of social play remain debated (Graham & Burghardt, 2010; Palagi, 2018). Notably, less research exists on the proximate mechanisms of social play (Martin & Caro, 1985; Spinka, Newberry & Bekoff, 2001; Held & Špinka, 2011; Pellis *et al*., 2023), with as exception the extensive neurobiological research of dyadic play in rats (Achterberg & Vanderschuren, 2023). Social play is a highly active social interaction, often characterized by fast exchanges of various behaviors with one or multiple participant(s) encompassing temporal patterns (Mielke & Carvalho, 2022). To achieve social play and to overcome the risk of play breaking down, potential proximate mechanisms are behavioral flexibility and the capacity to handle rapid exchanges (Held & Špinka, 2011), predicting other’s responses as well as potential unpredictability (Spinka *et al*., 2001; Palagi *et al*., 2016), or effective communication to avoid escalating a play fight into a real fight (Špinka, Palečková & Řeháková, 2016b). Play participants thus presumably rely on social skills with similar importance for cooperation, communication, or reciprocity (Wright *et al*., 2018).

Play signals are often proposed as key mechanism to facilitate social play (Bekoff, 1974). Play signals are communicative behaviors (e.g. facial expressions, limb movement, postures, or vocalizations) which function to regulate social play by conveying playful intentions, or by soliciting, maintaining, or modifying play (Bekoff, 1974; Yanagi & Berman, 2014a, 2014b). Successfully signaling the playful intent of a physical encounter, rather than a threatening one, is important to avoid misinterpretation and potentially costly outcomes (Bateson, 1955; Kuczaj & Horback, 2013; Pellis & Pellis 1996). In this context, such a meta-communicative signal changes the meaning of a subsequent behavior (Amici, Oña & Liebal, 2022). One way of providing partial support for this hypothesis is assessing whether signals mostly precede so-called contact offensive patterns (Maglieri *et al*., 2023), which are play behaviors involved during play fighting (e.g. wrestling, chasing, rough-and-tumble play). A more conclusive way to provide support is demonstrating the potential for misinterpretation, and thus the possibility of play escalating into fight, while demonstrating that the signals indeed reduce such escalation. This is for instance shown in Hanuman langurs (*Semnopithecus entellus*) which use ‘play-face’ (i.e. relaxed open-mouth display, van Hooff, 1972) as aggression-reducing signal (Špinka, Palečková & Řeháková, 2016a). Depending on the primate species, play-face might in addition, or only, reflect the sender’s internal motivation (Iki & Kutsukake, 2022) or maintain play bouts (Waller & Cherry 2012), among other proposed functions.

Research on play signals in primates has predominantly focused on play-face, but primates use other non-facial, bodily expressions that have been hypothesized to solicit, maintain, or modify play (Kipper & Todt, 2002; Palagi *et al*., 2016). Firstly, regarding the soliciting function of a play signal, research shows that rhesus monkeys (*Macaca mulatta*) use multiple postures to initiate play, with ‘leg-peeks’ leading to play initiations by the receivers of such expressions (Yanagi & Berman, 2014b). When comparing to matched-control data, further work found that young rhesus monkeys initiate play faster after a ‘crouch-and-stare’ than after no signal (Yanagi & Berman, 2014a). Also gorillas (*Gorilla gorilla gorilla*) start playing faster after a body signal compared to no signal (though note n = 2) (Weigel & Berman, 2018). Self-handicapping (i.e. a subject puts itself in a disadvantageous position that restricts the sender’s ability to tactically interact with another or easily defend themselves) is also proposed as signal to initiate social play, next to its function to convey playful intent (Spinka *et al*., 2001). For example, bonobos (*Pan paniscus*) are more likely to be engaged in social play after a period of solitary play which involved self-handicapping than when solitary play involved self-directed behaviors (Palagi, 2008). Secondly, a play signal may also maintain play. In line with evidence from research on play-face, rapid facial mimicry (i.e. an automatic, rapid mirroring response of a facial expression) during play is correlated with longer play bouts in chimpanzees (*Pan troglodytes*) (Waller & Dunbar, 2005) or geladas (*Theropithecus gelada*) (Mancini, Ferrari & Palagi, 2013). Recent work investigating the play initiating effects of face-to-face orientation in Japanese macaques (*Macaca fuscata*) shows that subsequential play bouts last longer compared to when its dyadic participants did not show face-to-face orientation before the start of the bout (Iki & Hasegawa, 2020). Thirdly, signals may also further modify the play dynamics. Japanese macaque dyads with face-to-face orientation before a play bout have both a more equal chance to gain advantage over their partner, compared to when only one partner faces the other (“unilateral gaze” as opposed to “mutual gaze”) (Iki & Hasegawa, 2020, 2021). Play dynamics also include differing levels of intensity and play types, and as such, signals may lead to more play fighting or wrestling, thereby building on the meta-communicative hypothesis. For instance, rhesus macaques (*Macaca mulatta*) initiate more intense play such as chase after using ‘crouch-and-stares’ signals (Yanagi & Berman, 2014b). In line with the meta-communicative hypothesis, switching or reversing roles between who signals and who subsequently initiates play may reveal a mutual understanding of the signaler’s communication. This type of turn-taking would in particular be interesting if the receiver of a signal becomes the one starting the play, as it would suggest that the signaler has successfully conveyed its intent (Yanagi & Berman, 2014b). If roles remain retained, and thus if the signaler acts mostly as play initiator, it could question the successful communication of playful intent and consequently, mutual understanding of the context.

Despite these various findings, the use of bodily expressions as signals, and their potential for a soliciting, maintaining, or modifying function for primate social play remains poorly understood. This is partially due to the lack of research on non-facial expressions, but another contribution concerns terminology and the applied analytical approach which varies between studies. This heterogeneity impacts data interpretation of the various functions of a signal. First, a re-occurring element is that the literature tends to label candidate signals as signals, without verifying their actual signaling and, thus, social effect on others and the interaction. The recommended practice is to apply unambiguous terminology, and this relates to expressions that are evidence-based signals versus expressions that are (merely) observed to be present during the interaction. A second important notion is the choice of analytical comparison. Social play may occur without the presence of a signal (Yanagi & Berman, 2014a) and, therefore, given that play is present, the research question should preferably focus on the potential of a candidate signal to change that subsequential play (i.e. through changing its probability, duration, etc.). Yet, current research on primate play signals tends to use two different analytical comparisons, each implying differing interpretations (Fig. 1).

**Figure 1:**
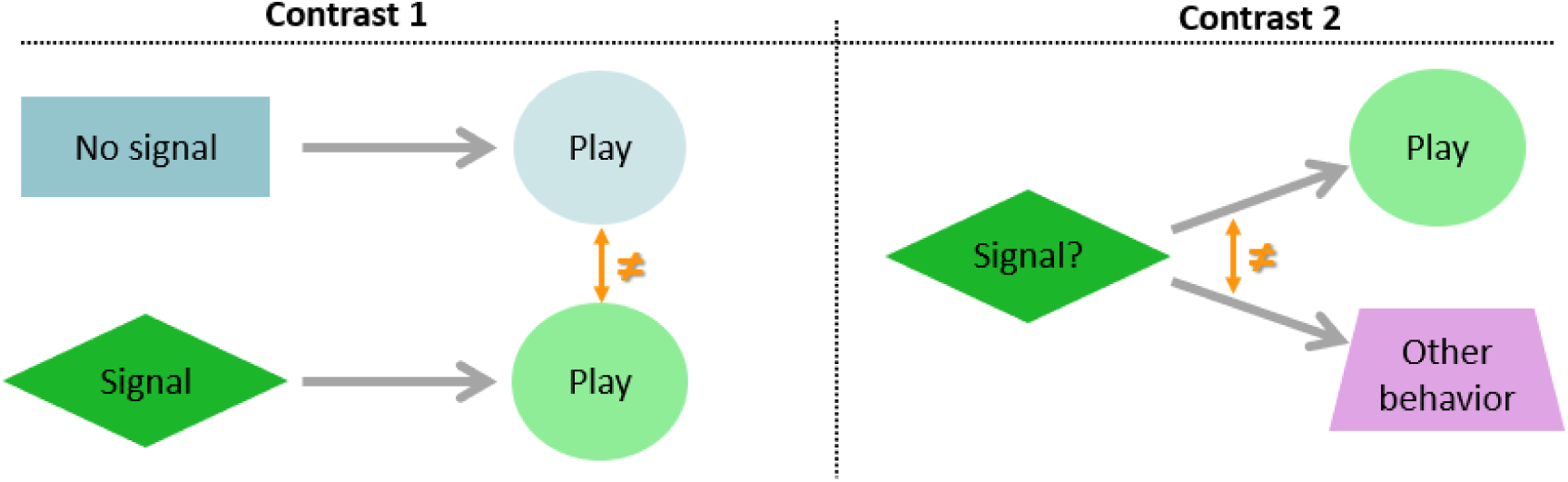
Contrast 1 tests whether the presence or absence of a signal will change the following play dynamics. Contrast 2 tests whether an expression will reliably lead to a specific action such as play, or other behavior. Contrast 1 aids in understanding the function of a signal, and Contrast 2 in understanding a candidate signal’s reliability.

The first question compares play that follows from a signal to play that does not follow from a signal (i.e. Contrast 1: signal to play vs. no signal to play). Contrast 1 tests whether the presence or absence of a signal leads to a difference in the subsequent play interaction. This comparison may for instance test whether a signal elicits play more than no signal (i.e. soliciting function) or whether play preceded by a signal lasts longer than when not preceded by a signal (i.e. maintaining function) (e.g. (Iki & Hasegawa, 2020)). The second question compares an expression resulting in play to the same expression resulting in other behavior (i.e. Contrast 2: signal to play vs signal to no play). Contrast 2 assumes, given that an expression is present, that it will (more or less) reliably lead to play or other behavior. This contrast thus provides additional information about the specificity of an expression. As parallel study topic, gesture research usually analyses the co-occurrence of specific expressions (e.g. limb movement) with social play versus other contexts. This research, therefore, tends to apply a Contrast 2 comparison, as questions may concern whether a gesture reliably conveys a specific meaning across contexts (Hobaiter & Byrne, 2014), or whether gestures are adjusted to the receiver’s identity (Fröhlich, Wittig & Pika, 2016). Note that the questions from both contrasts do not exclude one another, merely that Contrast 1 allows to understand if an expression is indeed a signal, whereas Contrast 2 leads to understanding the reliability of that expression. To conclude, to infer the function of a signal and how it may change the subsequent social play, a direct contrast of two conditions containing either a signal or not is required. Our study focuses on Contrast 1 as analytical comparison.

### Animal model

Common marmosets are cooperative breeders living in extended family groups (Digby & Barreto, 1993). Social play forms an important part of the common marmoset’s behavioral repertoire and all age classes show play throughout their lives (Stevenson & Poole, 1976; Voland, 1977). In particular, social play increases with immature age and parents are equally important play partners as peers (Godard, Burkart & Brügger, 2024). Research also shows evidence for an opiate mechanism, with morphine administration leading to increased social play in juveniles (Guard, Newman & Roberts, 2002). Importantly, social play is characterized by cooperating in space and time, with various participants expressing different play types, making choices on whom to engage with when, and potentially using signals to regulate all these aspects. To regulate group interactions, marmosets rely on particular communication and socio-cognitive skills (Burkart *et al*., 2022), and next to cooperatively carrying, provisioning, and protecting of infants (Erb & Porter, 2017), marmosets show extensive cooperation in other domains as well, such as coordinated territorial defense (Lazaro-Perea, 2001) or vigilance (Phaniraj, Brügger & Burkart, 2024). As such, social play forms an ideal candidate behavior to further investigate marmosets’ cooperative skills. Yet, how exactly the spatial and temporal aspects of social play are achieved in marmosets is not known. For instance, a 1981 study on play patterns found that marmosets play in longer play sessions consisting of multiple bouts, with chase usually occurring in the middle of a bout (Chalmers & Locke-Haydon, 1981). But overall less attention has been given to play in marmosets as a form of communication, and neither much research has been done on the use of signals to regulate play dynamics. Some propose that marmosets use signals as invitation to play (Voland, 1977), though detailed quantitative analyses are lacking, and others have dismissed the notion of signals on the basis that agonistic elements are absent and, thus, that there is no need for aggression-reducing signals (Stevenson & Poole, 1982). Nevertheless, play signals may have multiple functions (i.e. soliciting, maintaining, or modifying play), and so dismissal of signal presence in marmosets based on one function may not have sufficient theoretical grounding. Moreover, the extensive communicative skills of marmosets and their daily cooperative acts with others, renders the notion of marmosets using signals during play, or at most a form of gestural communication, a plausible idea (Burkart *et al*., 2022). An overall important point is that, despite the general assumption that social play should be easily recognized in other species, it is not necessarily a straightforward behavior to disseminate scientifically. Many experts on the topic emphasize this challenge, thereby highlighting the difficulty of the heterogeneous character of social play, including the presence of a long-standing belief that play is not biologically relevant (Graham & Burghardt, 2010). It is therefore not surprising that little research has been done on the use of signals during play in common marmosets, and their small body size and rapid movements provide additional difficulty for empirical research.

## PREDICTIONS

The main goal of this paper is to establish how body posture in marmosets is used as play signals, by directly comparing play resulting from the presence of a signal or not (Fig. 1). Concretely, we hypothesized that a signal would solicit play, maintain play, and modify play. Based on this we predicted that: (1) A signal increases the probability to play; (2) A signal extends the duration of play; (3) A signal leads to more diverse play; and (4) A signal increases the probability of intense play. Additionally, we explored the following questions based on general observations: (1) Do subjects take turns acting as sender or receiver?; and (2) Do signals get more dyads involved in the following play states?

## METHODS

### 1. Sample

The dataset used in this study comes from a larger data collection (Godard *et al*., 2024), to which author JECA contributed. The dataset concerns observational data from three family groups, with a total of 12 common marmosets (5 female). Each group consisted of two parents and two juveniles (see Supplemental Info, SI). All study subjects were captive-born, housed in family groups in temperature-controlled indoor enclosures (equipped with climbing material, platforms, branches, enrichment objects, sleeping boxes, and bark mulch as substrate) with regular access to an outdoor wire-meshed space (similar materials provided as indoor enclosures). Animals were fed twice per day (i.e. once vitamins and mash and once fresh fruit and vegetables) with an additional protein source and gum in the afternoon.

### 2. Data collection

Each group was videorecorded for a total of five (i.e. family 1 and 2) and four (i.e. family 3) sessions of 30 minutes each. For further behavioral coding the middle 15 minutes (07m30s -22m30s) from each session were used. As each session recorded all family subjects simultaneously, this resulted in 840 minutes of data (i.e. 15 minutes x 14 sessions x 4 subjects). To limit disrupting the naturally occurring play, we conducted recordings in the home enclosures. This resulted in various obstacles obscuring the view in the recordings and so we set up three Go Pro Hero 9 cameras to record from different angles (1: front of cage, 2: back of cage, 3: top of cage, see SI). This ensured a qualitative recording of the fast-paced and high-frequency behaviors of the small marmoset monkeys. All subjects were marked for individual identification, by differently located shaving markers on the tails. Data was collected between May 2021 and February 2022, either before, during, or after feeding time. During recording outdoor access was restricted and water was available *ad libitum*.

#### 3. Video coding

All 14 sessions were manually coded with a frame-by-frame (25fps) analysis and from three synchronized video angles, using the software INTERACT (Mangold GmbH). We applied focal sampling and thus each session was coded four times to cover all subjects per family group, with focus on: frequency and duration of signals and play types (Table 1, SI), and the initiating and receiving subject of the signal or play type. This resulted in approximately one week coding time per session. AO coded the majority of the data, with contribution from JECA, and a second observer independently coded 14% of the data, resulting in an excellent inter-observer reliability (Intraclass Correlation Coefficient, range: 0.98-1.00).

**Table 1:** Operational definitions.

| Term | Definition |
| --- | --- |
| **Play state** | A play state is a sequence of one or more play bout(s), which in turn may consist of one or multiple play type(s). A play state starts either with a signal or a play type. A state ends with a long break (i.e. rest) or a new signal, which marks the start of the next state. |
| **Play bout** | A play bout consists of a unique dyad performing either a play signal(s) or play type(s). A bout starts with the first signal or play type. A bout ends when: 1) one of the two participants initiates play with a new subject, resulting in a new dyad and a new bout; 2) both participants stop playing and a new dyad starts playing; or 3) when both participants stop playing for a long break. |
| **Play signal** | A signal is a posture during social play that occurs without physical contact and is non-essential to perform the actual play. The posture lasts for minimum 3 frames, is directed toward another subject including visual attention to the other. All signals were coded for frequency and duration. |
| **Play type** | A play type starts when a subject physically engages in a non-aggressive manner with another subject in any of the six play types. This is either with the first chase movement for *chase*, or the first body contact for the remaining types. A play type ends when subjects break physical contact or stop the *chase* movement. All types were coded for frequency and duration, except *touch* and *pounce* which do not have a duration. |
| **Breaks** | Breaks are either short (i.e. pause) or long breaks (i.e. rest). These are defined by a semi-supervised classifier. |
| **Rest** | Rest is a long break between states, and the break contains neither a signal nor play state. |

### 4. Ethogram

To compile the ethogram (Fig. 2, SI) we first assembled an overview of play behaviors and signals described in the literature (Voland, 1977; Chalmers & Locke-Haydon, 1981; Stevenson & Poole, 1982). Next, we had a broad viewing of the recordings, followed by group discussions to agree on the behaviors and their descriptions. A main question is whether an observed candidate signal is categorized as play signal or play type. Signals are mostly non-physical engagements with conspecifics, consisting of one or multiple modalities including olfactory, acoustic, or visual cues (Palagi, 2008). Our study focuses on the visual modality only, which includes various postures previously described as candidate signals of marmoset play. Play signals also often include visual attention to others (Yanagi & Berman, 2017). The start of a play type is described as the ‘first exchange of physical contact’ (e.g. (Iki & Hasegawa, 2020). This means that upon the first touch between participants a play bout officially starts, and the moment that physical contact breaks is the end of that play type, and potentially also bout if no other play type immediately follows. As such, a bout can consist of various play types within a dyad, and a play state can include multiple bouts interspersed with short breaks. Moreover, movements that are relevant or essential to the performance of play, such as grabbing the other before a wrestle, are considered to be part of the play bout and should not be labelled as play signal (e.g. Pellis & Pellis, 1996). The resulting ethogram includes three play signals: hide, stalk, and supine, of which only hide has not yet been described in the marmoset literature; and six play types, namely pounce, pull, touch, wrestle, chase, and catch. Supine is also called a self-handicapping posture (Petrů *et al*., 2009; Fröhlich *et al*., 2016). Wrestle is often labelled rough-and-tumble play (Birnie *et al*., 2012), though this may or may not include both wrestle and chase, so for sake of clarity our ethogram separates the two types. Previous research mentions ‘looking-between-legs’ in marmosets (Voland, 1977), but we did not observe this in any of our recordings.

**Figure 2:**
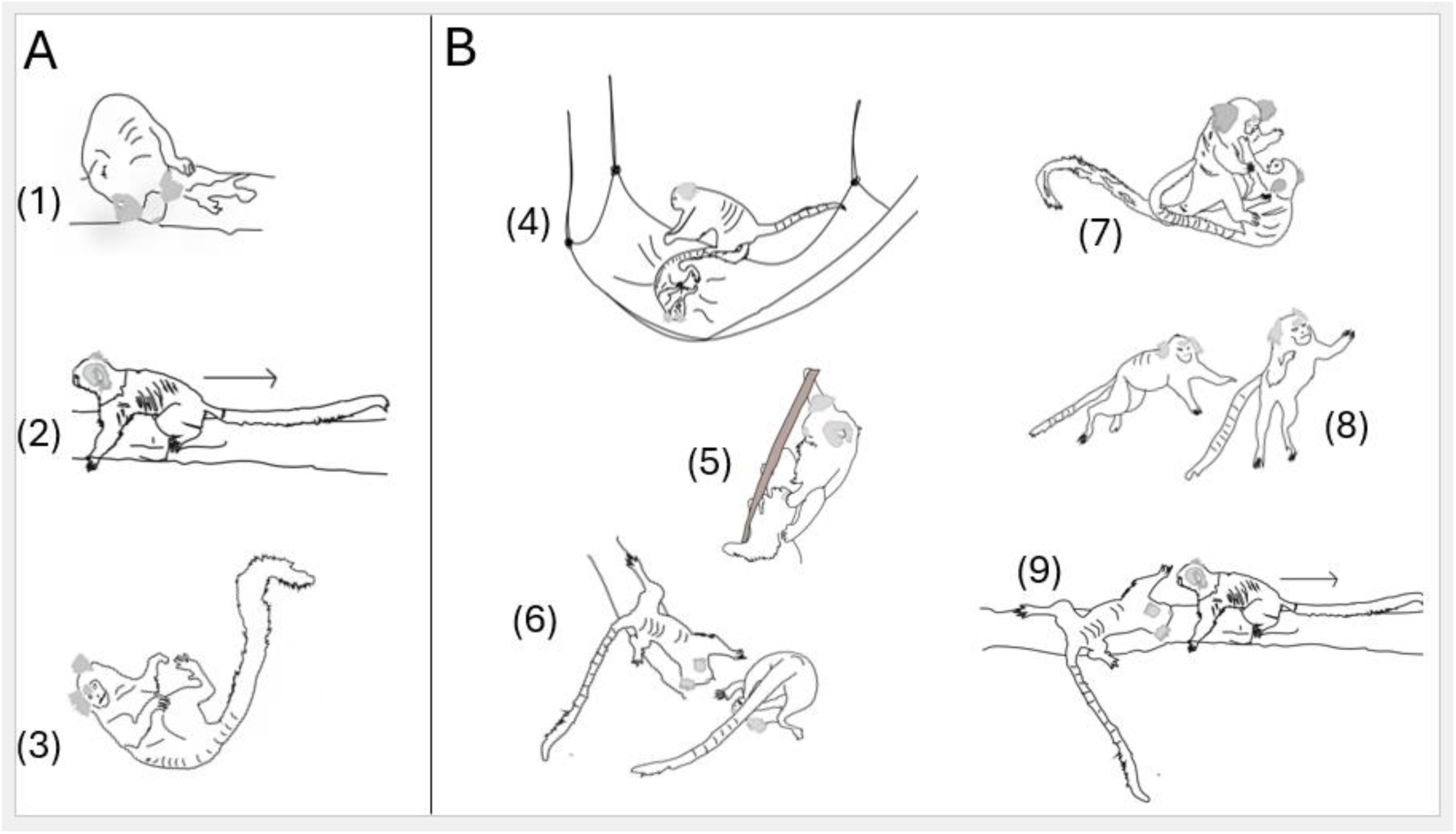
Illustrated ethogram (Godard *et al*., 2024). **A** shows 3 signals: (1) Hide, (2) Stalk, (3) Supine; **B** shows 6 play types: (4) Pounce, (5) Pull, (6) Touch, (7) Wrestle, (8) Chase, and (9) Catch.

### 5. Data analysis

In the first part of our analysis, we used a Bayesian multi-state time-to-event model (Andersen, Abildstrom & Rosthøj, 2002; Asher *et al*., 2017; Luo *et al*., 2021). Specifically, we model the continuous-time transitions between three behavioral states: a signal state, a play state, and a rest state (Table 1). Behavioral states were defined at the level of the entire family group rather than at the individual level, due to strong dependencies between the behavior of individuals in these small groups (e.g. if subject A is playing with subject B, this constrains the behavior of subject C, who may not have an eligible partner to play with). These dependencies arise from both the captive housing and the social system of the marmosets. Analyzing group-level behavior (e.g., is there any play going on in the family group at time point *t*?) overcomes this problem and makes the statistical model tractable.

A general challenge for analyzing naturalistic behavioral sequences is how to process the lags, or latencies, between behavioral states. For example, if there is a 5 second lag of no activity between two play behaviors, should this be treated as a distinct rest state, or rather a continuous play state punctuated by a pause? Many authors use somewhat arbitrary cutoffs for this determination, such as 10 seconds between behaviors. But there can be ambiguous lags near the boundaries of these cutoffs, introducing bias in the construction of sequences. To avoid these pitfalls, we utilized a data-driven mixture model that probabilistically classifies lags as either short pauses or longer rests. The resulting classifications allowed us to segment the behavioral sequences appropriately for use in subsequent analyses.

When analyzing the duration of play states in subsequent analyses, we used a multilevel model with a Gamma likelihood, which accommodates potential deviations from the assumption of exponentially distributed waiting times. When analyzing the counts of play types and the number of dyads within a play state, and the analysis of how many dyads participate in a given play state, we used an ordinal likelihood. For the analysis of the presence/absence of specific behaviors within play states, we used a Bernoulli likelihood. For the analysis of turn taking in who is the sender/receiver, we used a Categorical likelihood reflecting the three possible responses (roles reversed, roles retained, other).

Session-level random effects were included in all models to account for the non-independence of behaviors within a given recording. In the results section, we report the average effects/predictions, marginalizing over session and group-level differences. Analyses were run in R (version 4.3.2 R Core Team, 2015) using the RStan package (Stan Development Team, 2020), which fits Bayesian models using Hamiltonian Markov chain Monte Carlo (MCMC), assessed using standard diagnostics (number of effective samples, the Gelman–Rubin diagnostic, and visual inspection of trace plots). We give an in-depth, formal definition of our models, as well as several diagnostics of model performance, in the supplementary materials.

When reporting the estimates from our models, we give posterior means, 90% highest posterior density intervals (HPDIs), and the posterior probability of direction (pd), when assessing a directional effect. The pd is a continuous measure of the strength of evidence, but to self-enforce consistency we refer to pd> .90 as “strong” evidence, pd > .80 as “moderate”, and pd <= 0.8 as “weak”.

## RESULTS

### Lag classification

The data-driven mixture model classified time lags between states as either short pauses or longer rests (Fig. 3a). Using this outcome, we find that marmoset play occurs as play states which consist of one or multiple bouts in rapid succession (Fig. 3b).

**Figure 3:**
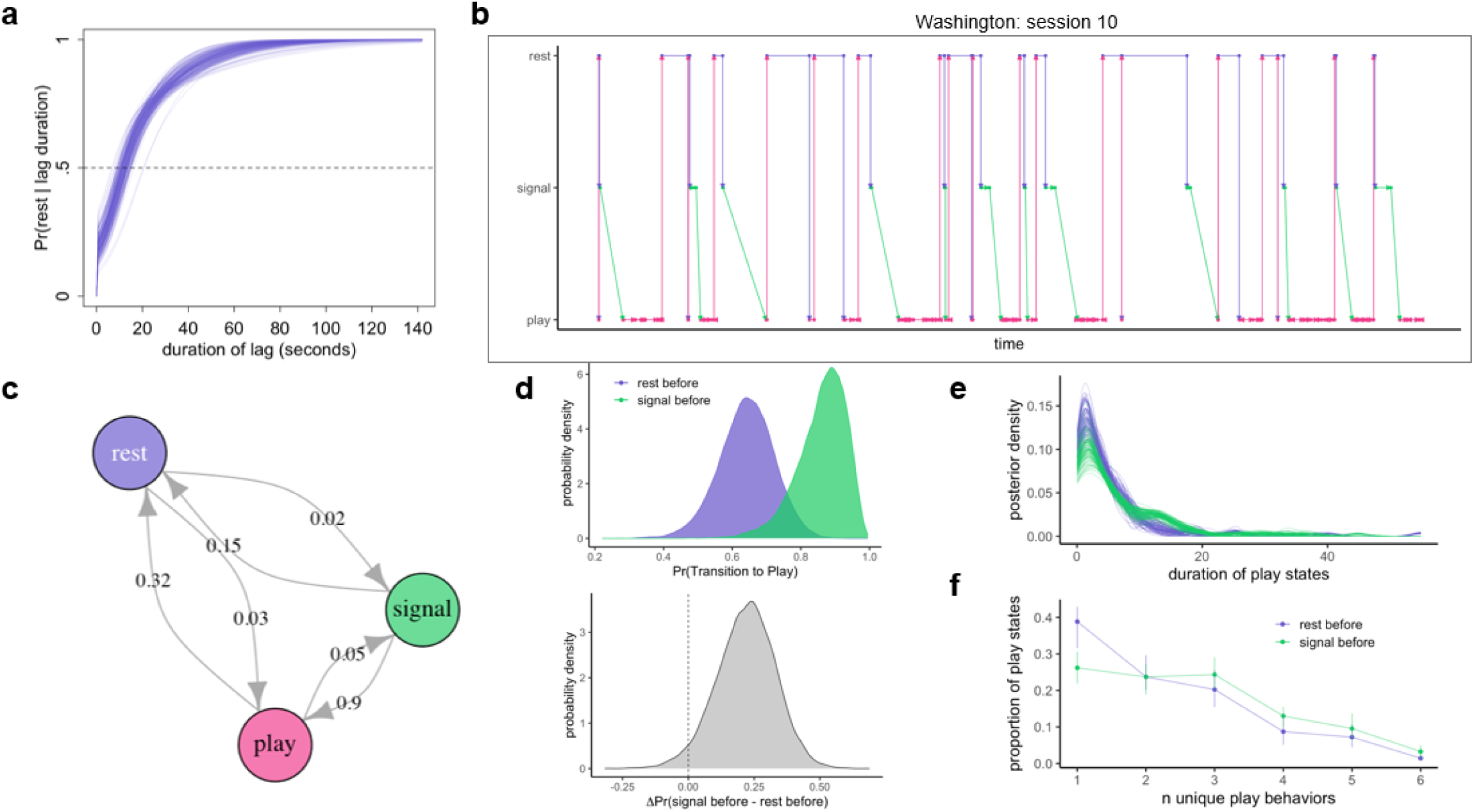
**(a)** Probabilistic classification of lags between behavior as a rest state (i.e. long break) or pause (i.e. short break), as a function of duration. **(b)** Posterior mean behavioral sequences for a single session as example, with rest states inserted based on the classifier shown in panel a. **(c)** Posterior mean transition rates between each state, marginalizing over sessions. **(d)** Posterior probability of transitioning to play from either rest or signal (top) and the difference in those two probabilities (bottom). **(e)** Posterior distribution of the duration of play states depending on whether it was preceded by rest or signal. **(f)** The number of unique behaviors within the play state, with points representing posterior means and bars for 90% HPDIs. Pink: play states; Green: signal states; Purple: rest states.

### Descriptive data of overall play

All reported descriptive values are from the multi-state model classifier, adjusted for uncertainty and 90% CI. In total there were 166 [150, 184] play states with an average duration of 5.47 [4.91, 5.95] seconds. Most play states included chase (58.5% [55.3%, 61.3%]) and wrestle (56.6% [53.9%, 60%]), followed by pounce (27.4% [24.9%, 30.6%]), touch (28.8% [26.5%, 30.8%), pull (14.4% [13%,15.8%]), and catch (1.87% [1.63%, 2.16%]). Most play dyads consisted of the juvenile twins playing with each other (57.5%), with the remaining dyads involving the parents (42.5%). Out of all play states, 45.8% [41.4%, 53.2%] were preceded by at least one play signal, and for these states, signals occurred on average 11.38 [9.025, 13.703] seconds before a play state (i.e. measured from the end of the signal to the start of the play). The other play states started without signal and thus immediately with a play type. Signals were hide (32.2% [31.1%, 32.9%] of signal states), stalk (38.4% [37.4%, 39.5%] of signal states), and supine (50% [48.9%, 51.2%,] of signal states).

### Prediction 1: A signal increases the probability to play

The average probability of transitioning from a rest state to play state is .641 [.512, .774], while the probability of transitioning from signal to play is .860 [.753, .976]. The difference in these two probabilities offers strong evidence that play is more likely to occur after signaling Pr(Play|Signal) – Pr(Play|Rest) = .219 [.032, .398], pd = .968 (Fig. 3d). Alternatively, the odds of transitioning from signal to play are about 5 times higher than to start playing after rest (OR = 5.266, 90% CI = .516 –10.353).

### Prediction 2: A signal extends the duration of play

The average duration of a play state that was preceded by rest was 3.27 [2.19, 4.29] seconds, while play states preceded by a signal had an average duration of 4.08 [2.48, 5.57] seconds. Play states preceded by a signal lasted an average of 0.81 [-.76, 2.36] seconds longer than those preceded by rest, with the evidence being moderate (pd = .808) (Fig. 3e).

### Prediction 3: A signal leads to more diverse play

The average number of unique play behaviors during a play state that was preceded by rest was 2.35 [2.05, 2.66], while the average number of play behaviors during a state preceded by a signal was 2.72 [2.34, 3.10]. Play states preceded by a signal had an average of 0.36 [-.09, .83] more unique behaviors than states preceded by rest (pd = .903). To explore whether this increase could be explained by the longer duration of play states preceded by a signal (Prediction 2), we fit an additional model that included the duration of play as a prediction. This adjustment for duration then changes the question to: “does the *rate* (as opposed to count) of unique play behaviors increase when preceded by a signal?”. We found that, after adjusting for duration, play states after a signal had an average of .22 [-.17, .62] more unique play behaviors than play states preceded by rest, with moderate support (pd = .82) (Fig. 3f).

### Prediction 4: A signal increases the probability of more intense play

The average probability of wrestling during a play state preceded by rest was .50 [.40, .61], while the average probability of wrestling following a signal was .64 [.52, .76]. Play states preceded by a signal had an average of .13 [-0.02, .28] greater probability of including wrestling than play preceded by rest (pd = .922), thus providing strong support for this prediction (Fig.4). The average probability of chase during a play state preceded by rest was .58 [.48, 69], while the average probability of chase following a signal was .58 [.46, .69]. Play states preceded by a signal had an average of -0.01 [-.15, 0.14] less probability of including chase than states preceded by rest, thus providing weak support for this prediction (pd = .539) (Fig. 4).

**Figure 4:**
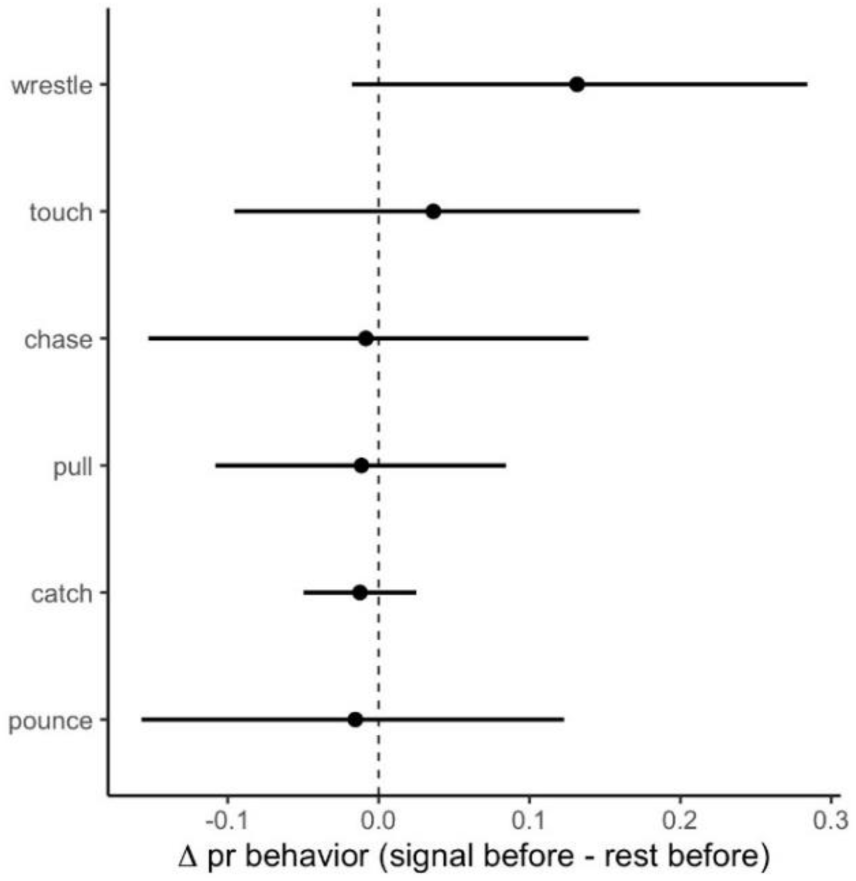
Average difference in the probability that a play state contains a behavior when the play was preceded by a signal compared to a rest state. Points represent posterior means and error bars represent 90% highest posterior density intervals.

### Exploratory question 1: Do subjects take turns acting as sender or receiver?

We asked whether the sender of a signal (i.e. signaler) became the receiver of the following play state, which would imply that the receiver of the signal becomes the initiator of the play (i.e., role reversal or ‘turn-taking’). We found that the average probability of a role reversal between signal and play state was .295 [.194, .392], while the probability of retaining roles was .355 [.250, .461]. The remaining probability of .350 [,.255, .445] is assigned to cases in which the initiator of play was neither the signaler nor receiver of the signal (or vice-versa) (i.e. another dyad initiates play). When analyzing role reversal among the same dyads, we find that the probabilities of either retaining or reversing roles between signaling and play are very similar (Pr(reversal) – Pr(retain) = -.061 [-.243, .121]).

### Exploratory question 2: Do signals get more dyads involved?

In play states preceded by rest, the average number of dyads is 1.27 [1.13, 1.40], and when play is preceded by signal, the average number of dyads is 1.34 [1.14, 1.53]. The difference (signal – rest) is 0.068 [-.151, .280], pd = .691, providing weak evidence that signals lead to more unique dyads playing.

## DISCUSSION

### Main results

The main goal of this paper was to investigate how body signals in marmosets are used as play signals, by directly comparing play preceded by a signal or not. We applied a Bayesian statistical approach with a multi-state time-to-event model to test four predictions and two exploratory questions.

First, we found strong support that a signal increases the probability to play, which provides evidence for a soliciting function of signals during marmoset play. In addition to existing research in for instance bonobos or rhesus monkeys (Palagi, 2008; Yanagi & Berman, 2014a), we now provide evidence that common marmosets also use signals to initiate play. Important to our research question, we found that when signals were present, the odds were about 5 times higher for marmosets to start playing.

Second, we found moderate support that a signal extends the duration of the following play, which provides evidence for a maintaining function. This result is in line with research in rhesus and Japanese macaques (Wright *et al*., 2018; Iki & Hasegawa, 2020). Some researchers argue that longer play states, due to a preceding signal, are evidence of those signals serving a meta-communicative function. The idea is that signals avoid escalation into aggression, and thus a premature termination of that play, and therefore lead to longer play interactions. Yet, we follow the notion that to provide evidence of such aggression-reducing function, research should demonstrate observations of the presence of such escalations, and then further make direct comparisons between the presence and absence of signals in relation to the presence or absence of escalation and the duration of the following play (Špinka *et al*., 2016b). In this study we did not observe any aggression escalation during social play, and thus we cannot use the duration results to interpret a potential meta-communicative function. Similar to our design, recent work in Japanese macaques compared the effect of the presence versus absence of play-face on the following play (Iki & Kutsukake, 2022). The researchers found that play indeed lasted longer after play-face, but they found no evidence that play-face avoided escalation or engaged reluctant others. Rather, play-face was interpreted as indicating internal motivation to play, and perhaps a similar interpretation can be applied to the marmoset signals. Research often suggests that play motivation stems from a playful mood (i.e. a long-term affective state) (Mendl, 2010). An interesting avenue to understand if body signals reflect play motivation, and thereby prolong subsequential play, is to investigate the underlying affective state during play (Ahloy-Dallaire, Espinosa & Mason, 2018; Adriaense *et al*., 2020).

Third, we found moderate support that a signal leads to more diverse play, even when controlling for duration effects, thus providing evidence for a modifying function. Yet, which exact mechanism drives this result is unclear. The literature suggests that a signal might establish a mutual understanding that the current situation is indeed play, and for instance not a risky context. This may lead to subjects uninhibitedly exploring the play interactions with others and allowing a richer context, thereby resulting in more unique play types. Further research would be required to understand this finding.

Finally, also in line with the modifying hypothesis, we found strong support that a signal leads to more wrestle, with the support for chase being weak. In other words, marmosets that preceded their play with a signal were more likely to wrestle during following play, than when play was not preceded by a signal. Wrestle is frequently used in primate research as indicator for intense play or play fighting, meaning that this type has a higher chance of escalation into a real fight. Our result implies that signals convey playful intent and a shared understanding that wrestle is only play and not a real fight, which then allow play fight to occur more when a signal has been given. This finding supports a meta-communicative function of signals (Maglieri *et al*., 2023), and as common marmosets in general show both within-and between-group aggression, they may indeed require signals to avoid these situations in a play context. Overall, the finding that signals lead to more wrestle forms an interesting addition to the meta-communicative hypothesis, yet, given that we did not observe any aggression during play in our dataset and thus cannot directly test for an aggression-reducing effect (see argument above), more research is required for a more conclusive interpretation. Regardless, the finding of a signal leading more to wrestle, with strong support from the statistical model, provides further convincing evidence for a modifying function of play signals in marmosets.

When exploring the data further, we found close to no difference in probabilities of subjects retaining or reversing their role of sender or receiver. In other words, if subjects were acting as the signaler, there was about 50% probability they became receiver of the following play (i.e. role reversal or turn-taking) or the initiator of that play (i.e. role retainment). Comparably, rhesus macaques who are receivers of a signal initiated about 63% of the following play (Yanagi & Berman, 2014b). Though our results cannot provide conclusive evidence for complete turn-taking, the similar probability to retain or reverse role does indicate that in about half of the time subjects take turns in who is signaling and who is initiating the play interaction. This implies that marmoset signalers do not necessarily drive the entire play interaction, and that signal receivers potentially have a mutual understanding of the signaler’s communication. Such turn-taking, and in particular a receiver of a signal becoming the initiator of the following play, may suggest that these signals successfully convey the intent of the signaler (Townsend *et al*., 2017; Weigel & Berman, 2018). Common marmosets are known for their turn-taking skills (Burkart *et al*., 2022), such as vocal turn-taking (Takahashi, Narayanan & Ghazanfar, 2013), motor coordination in a joint action task (Miss & Burkart, 2018), and vigilance coordination during feeding (Phaniraj *et al*., 2024). Our study therefore contributes to further understanding the extent of turn-taking in marmosets. Additionally, we find that 43.5% of the dyads concern play between a juvenile and a parent, which supports previous findings of the extensive involvement of marmosets parents in social play (Stevenson & Poole, 1982; Godard *et al*., 2024). An important question arising is whether it is mostly the juveniles that drive the entire play interaction, with parents giving in to the signaling and play initiations of the juveniles (i.e. role retainment), or whether parents also take a leading role (i.e. role reversal). Descriptively we find that within juvenile-parent dyads 25% role reversal occurs, and though roles are thus mostly retained, to some extent parents also drive the entire play interaction with their juveniles.

Finally, we explored whether signaling within a single dyad would result in multiple other dyads involved in playing afterwards, but we found weak support for this idea. This question arose from the interest in contagion (Adriaense *et al*., 2021), as play is suggested to have contagious effects on others and thereby spread a playful mood leading to multiple others playing as well. In a similar vein, it could be that visually signaling is picked up by others and thereby motivating them to play, as marmosets have the ability to extract information from social interactions of others (Brügger, Willems & Burkart, 2021). Our findings are in contrast with such interpretation, and we find no effects of signaling on others outside the signaling dyad. Yet, these findings support the interpretation that signals are communicative acts intentionally directed to another specific subject. Intentional use of a signal refers to the signaler’s visual attention toward the other, by visually monitoring them (Fröhlich & Hobaiter, 2018). Further detailed analysis of visual attention is required to investigate this interpretation.

To conclude, this study is the first to investigate how common marmosets use signals during play, in which we find strong evidence for body posture functioning as signal to solicit play in others as well as to lead to intense play in the form of wrestling. We also find moderate evidence that body posture leads to longer and more diverse play. Additionally, marmosets reverse their roles of acting as sender or receiver of signaling and subsequent play in approximately half of the time, suggesting a form of turn-taking. Taken together, these results imply that marmosets, through use of signals, share a mutual understanding of the playful context, thereby providing opportunity to not just start playing, but also to play longer, explore more, and play more intensely. To disentangle effects of sharing such understanding with others from, for instance, signals reflecting internal states such as play motivation, further detailed research is required on each of the functions.

### Statistical contribution

Social play is a compilation of fast-paced and varying behaviors, and to analyze this, statistical models ideally reflect these complex patterns. Yet, with many of the current methods, it remains difficult to investigate how candidate expressions indeed function as signal during primate social play. In particular, disentangling the different functions of signals often remains ambiguous. Other researchers have pointed out the need for advanced statistical methods (Wright *et al*., 2018) and our study aimed at meeting these requirements by applying a unique statistical approach.

First, we used a data-driven approach to classify play and thereby avoid arbitrary, and ultimately subjective, bout intervals. The use of such arbitrary intervals is frequent in current play research, with researchers usually conforming within a given species, such as 5 seconds for common marmosets (Mustoe *et al*., 2014) and rhesus macaques (Yanagi & Berman, 2017), 10 seconds for Tibetan and Japanese macaques (Wright *et al*., 2018; Iki & Hasegawa, 2020), or 2 minutes for chimpanzees (Heesen *et al*., 2021; van Boekholt, Wilkinson & Pika, 2024). Bout intervals will differ greatly between primate species due to their body size or agility, but also due to other contextual effects such as a (captive) environment, the total space available to play, the number of conspecifics present to become play mates, or even the type of play. One solution is to conduct a bout analysis of the observed dataset and apply the most common or smallest interval as criterion (Chalmers & Locke-Haydon, 1981), but this would still exclude intervals or ignore contextual effects. To account for this, our methods include a semi-supervised classification of time lags between behaviors as either short or long transitions. This resulted in an appropriate segmentation of the behavioral sequences, and thus gave a more accurate portrayal of the naturally occurring behavior. The resulting categorization was then used to investigate the key questions regarding signaling function.

Second, an ideal model would include all potential states of a play interaction, such as a play state, signal state, as well as a rest state in which no play or no signal occurs, which serves as baseline. To account for a baseline, an often-used method in primate research is matched-control sampling (MC): on day 1 behavior is measured for a certain period after a signal, and then a matched recording is done on day 2, with the same individuals, group, and context as day 1, but without a signal. Then analysis compares the two periods to examine how a signal may affect play. This MC procedure ensures a similar distribution of the data between the test and control day, yet, naturally occurring behavior is noisy, and potential effects on the behavior in day 2 might be due to unaccounted variables. Moreover, it is a known difficulty to find an exact match between two days. As an alternative, the multi-state model used in this study includes a rest state, which occurs within the same observation period of the signal and play state, and thus avoids the need to artificially create no-signal events at another timepoint.

A third characteristic is that a model reflects the delayed or accumulated effect of signal(s) rather than merely measuring immediate effects. Interesting approaches to study play include sequence based analyses (Mielke & Carvalho, 2022) though they lack a temporal component. Adding temporal information to a model takes not only into account which signal and play behaviors follow each other immediately, but also observed delays between signal and play (Chalmers & Locke-Haydon, 1981), as well as providing transition probabilities between states. The statistical approach of this paper is a multi-state time-to-event model, including both event and temporal information, which to our knowledge has not yet been used for animal play research.

### Future outlook

Some of the findings of this study make way for interesting future research. As such, this study coded the senders of a signal based on the sender’s visual orientation to another subject, who was then labelled the receiver. But it is difficult to strictly define how this visual contact was established, for instance through mutual or unilateral orientation, and research shows that depending on this orientation, different play dynamics may occur (Iki & Hasegawa, 2020). Future research with fine-grained measures of visual attention would benefit our understanding of the precise mechanisms of signaling and, in addition, would help to clarify the intentional use of such signal (Fröhlich & Hobaiter, 2018). Further, the supine signal is a self-handicapping posture, as also observed in for instance chimpanzees (Fröhlich *et al*., 2016). Self-handicapping postures restrict the sender’s ability to interact with others and it is suggested that this may train subjects for the unexpected (Spinka *et al*., 2001). The dataset in this study is not sufficient to test for this hypothesis, nor to disentangle supine specific effects from other signals on immediate play, but a more in-depth analysis of self-handicapping in common marmosets would be an interesting future direction. Finally, the manual coding of this dataset was undoubtedly time consuming, specifically due to the information on both event and temporal data. Future work involving algorithm detection of socially interactive behaviors with multiple subjects in noisy environments would be a great benefit to this research topic (Wiltshire *et al*., 2023), which would in turn allow a greater sample size and dataset for more extensive analyses of the signals.

Further, though signals clearly increase the probability to play, this study also shows that signals are not a necessary requirement to play, as in about 54% of play events there were no preceding signals. Comparison of the signal frequency to the frequency in other primates is difficult, as research varies in the modalities it includes to denote a play signal. For instance, signals may include both body signals and play-face expressions, with rhesus macaques showing such signals in 49% of play events (with 80% of these signals being play-face) (Yanagi & Berman, 2014b), signals may solely refer to face-to-face orientation, with Japanese macaques using this signal in about 80% before playing (Iki & Hasegawa, 2021), or signals may refer to a summary of multiple possible modalities including vocalizations, facial expressions, gestures, and tactile cues, with chimpanzees using these signals 69% of the playing time and bonobos 90% of the time (Heesen *et al*., 2021). Common marmosets have an extensive vocal repertoire and recent work such as CalliFACS (Correia-Caeiro *et al*., 2022) reveals the potential for intricate analyses of marmoset play-face (de Boer *et al*., 2013). Unfortunately an additional investigation of these modalities was not possible, as observations were done in the home enclosures with various objects blocking the (micro) view of play-face, and the nearby presence of other families hindering acoustic recordings. This study therefore focused on body signals as singular modality, yet future work on marmoset play should aim to apply a multimodal approach.

Next to future research, the study of play signals can be conceptually linked to other topics such as joint action. Recent work has proposed an interaction framework approach to measure joint actions by means of quantifying gestures or ‘coordination efforts’ (Genty *et al*., 2020). The approach suggests an entry phase, in which the actors initiate the joint action, followed by the joint action itself, and ends with an exit phase in which again expressions communicate the closure of an action (Baehren, 2022). Social play, being a coordinated interaction, is considered a joint action (Heesen *et al*., 2017), and so the conceptual overlap with the joint action framework lies in the use of signals to solicit and/or end social play, and whether play follows predictive and coordinated sequences that would imply the entry-to-exit framework idea (Holler, 2022). In this regard, our study contributes to understanding joint action in common marmosets, as we find that marmosets use posture as a signal to solicit play (i.e. entry phase) and to change play dynamics during play by having more diverse and intense play (i.e. middle phase), and marmosets also show role reversal of initiating play (i.e. necessary coordination during the framework) (Miss, Adriaense & Burkart, 2022). Marmosets do not seem to show exit phase expressions during play, though a multimodal approach and a fine-grained analysis of their visual attention could be beneficial in this regard. Importantly, the distinction with animal joint action research is that it usually assesses the presence of expressions throughout an interaction, whereas this study assesses the (consequential) function of those expressions on the interaction.

## CONCLUSION

To conclude, our study forms a twofold contribution by providing empirical evidence that common marmosets use signals multi-functionally during play and by applying statistical state-of-the-art methods to understand primate play. The main goal was to study postures as candidate signals of social play and we found evidence of such signaling function, as marmosets use postures to solicit, maintain, and modify play. We also found a form of turn-taking during signaling and play. These results imply that postures act as communicative signals in marmosets which may also convey mutual understanding and intentions. Our data-driven statistical approach modelled naturally occurring play events and their timing, avoided arbitrary event intervals, acknowledged group-dependency effects, and prevented the use of artificial control contexts. Overall, given the conceptual links between social play and cooperation and communication, this study on marmoset play provides a valuable contribution to further understanding these topics.

## Supporting information

Supplemental Stats

## Acknowledgements

We thank Hidir Sengül and Dominique Ziegler for animal care-taking and support during data collection. We also thank Rahel Brügger and Alice Godard for help with data collection, finalization of the play ethogram, and the generous sharing of the illustrated ethogram. This project received funding from the NCCR Evolving Language, Swiss National Science Foundation Agreement no. 51NF40_180888 (to J.E.C.A. and J.M.B.).

## Data Availability Statement

All data generated in the study and the code for analyses (R-script) will be made open access upon publication.

## Ethics Statement

All experiments were carried out in accordance with the Swiss legislation and licensed by the Kantonales Veterinäramt Zürich (license number: ZH232/19). All the procedures were fully non-invasive (degree of severity: 0). No recordings were conducted outside the home enclosure and no animal was food or water deprived during any point of this study.

## REFERENCES

Achterberg, E.J.M. & Vanderschuren, L.J.M.J. (2023) The neurobiology of social play behaviour: past, present and future. Neuroscience & Biobehavioral Reviews, 105319.

Adriaense, J.E.C., Koski, S.E., Huber, L. & Lamm, C. (2020) Challenges in the comparative study of empathy and related phenomena in animals. Neuroscience & Biobehavioral Reviews 112, 62–82.

Adriaense, J.E.C., Šlipogor, V., Hintze, S., Marshall, L., Lamm, C. & Bugnyar, T. (2021) Watching others in a positive state does not induce optimism bias in common marmosets (Callithrix jacchus), but leads to behaviour indicative of competition. Animal Cognition 24, 1039–1056.

Ahloy-Dallaire, J., Espinosa, J. & Mason, G. (2018) Play and optimal welfare: Does play indicate the presence of positive affective states? Behavioural Processes 156, 3–15.

Amici, F., Oña, L. & Liebal, K. (2022) Compositionality in Primate Gestural Communication and Multicomponent Signal Displays. International Journal of Primatology.

Andersen, P.K., Abildstrom, S.Z. & Rosthøj, S. (2002) Competing risks as a multi-state model. Statistical Methods in Medical Research 11, 203–215.

Asensio, N., Zandonà, E., Dunn, J.C. & Cristóbal-Azkarate, J. (2022) Socioecological correlates of social play in adult mantled howler monkeys. Animal Behaviour 186, 219–229.

Asher, L., Harvey, N.D., Green, M. & England, G.C.W. (2017) Application of Survival Analysis and Multistate Modeling to Understand Animal Behavior: Examples from Guide Dogs. Frontiers in Veterinary Science 4. Frontiers.

Baehren, L. (2022) Saying “goodbye” to the conundrum of leave-taking: a cross-disciplinary review. Humanities and Social Sciences Communications 9, 46.

Bekoff, M. (1974) Social Play and Play-Soliciting by Infant Canids. American Zoologist 14, 323–340.

Birnie, A.K., Hendricks, S.E., Smith, A.S., Milam, R. & French, J.A. (2012) Maternal gestational androgens are associated with decreased juvenile play in white-faced marmosets (*Callithrix geoffroyi*). Hormones and Behavior 62, 136–145.

Van Boekholt, B., Wilkinson, R. & Pika, S. (2024) Bodies at play: the role of intercorporeality and bodily affordances in coordinating social play in chimpanzees in the wild. Frontiers in Psychology 14.

De Boer, R.A., Overduin-De Vries, A.M., Louwerse, A.L. & Sterck, E.H.M. (2013) The behavioral context of visual displays in common marmosets (Callithrix jacchus). American Journal of Primatology 75, 1084–1095.

Brügger, R.K., Willems, E.P. & Burkart, J.M. (2021) Do marmosets understand others’ conversations? A thermography approach. Science Advances 7, eabc8790. American Association for the Advancement of Science.

Burkart, J.M., Adriaense, J.E.C., Brügger, R.K., Miss, F.M., Wierucka, K. & Van Schaik, C.P. (2022) A convergent interaction engine: vocal communication among marmoset monkeys. Philosophical Transactions of the Royal Society B: Biological Sciences 377, 20210098.

Chalmers, N.R. & Locke-Haydon, J. (1981) Temporal patterns of play bouts in captive common marmosets.

Ciani, F., DALL’olio, S., Stanyon, R. & Palagi, E. (2012) Social tolerance and adult play in macaque societies: a comparison with different human cultures. Animal Behaviour 84, 1313–1322.

Correia-Caeiro, C., Burrows, A., Wilson, D.A., Abdelrahman, A. & Miyabe-Nishiwaki, T. (2022) Callifacs: The common marmoset Facial Action Coding System. Plos One 17, e0266442. Public Library of Science.

Digby, L.J. & Barreto, C.E. (1993) Social Organization in a Wild Population of Callithrix jacchus. Brill.

Erb, W.M. & Porter, L.M. (2017) Mother’s little helpers: What we know (and don’t know) about cooperative infant care in callitrichines. *Evolutionary Anthropology: Issues*, News, and Reviews 26, 25–37.

Fagen, R. (1982) Skill and flexibility in animal play behavior. Behavioral and Brain Sciences 5, 162– 162.

Fröhlich, M. & Hobaiter, C. (2018) The development of gestural communication in great apes. Behavioral Ecology and Sociobiology 72, 194.

Fröhlich, M., Wittig, R.M. & Pika, S. (2016) Play-solicitation gestures in chimpanzees in the wild: flexible adjustment to social circumstances and individual matrices. Royal Society Open Science 3, 160278. Royal Society.

Genty, E., Heesen, R., Guéry, J.-P., Rossano, F., Zuberbühler, K. & Bangerter, A. (2020) How apes get into and out of joint actions: Shared intentionality as an interactional achievement. Interaction Studies. Social Behaviour and Communication in Biological and Artificial Systems 21, 353–386.

Godard, A.M., Burkart, J.M. & Brügger, R.K. (2024) The ontogeny of play in a highly cooperative monkey, the common marmoset. bioRxiv. https://www.biorxiv.org/content/10.1101/2024.05.28.595935v1 [accessed 4 August 2024].

Graham, K.L. & Burghardt, G.M. (2010) Current Perspectives on the Biological Study of Play: Signs of Progress. The Quarterly Review of Biology 85, 393–418.

Guard, H.J., Newman, J.D. & Roberts, R.L. (2002) Morphine administration selectively facilitates social play in common marmosets*†. Developmental Psychobiology 41, 37–49.

Heesen, R., Bangerter, A., Zuberbühler, K., Iglesias, K., Neumann, C., Pajot, A., Perrenoud, L., Guéry, J.-P., Rossano, F. & Genty, E. (2021) Assessing joint commitment as a process in great apes. iScience 24, 102872.

Heesen, R., Genty, E., Rossano, F., Zuberbühler, K. & Bangerter, A. (2017) Social play as joint action: A framework to study the evolution of shared intentionality as an interactional achievement. Learning & Behavior 45, 390–405.

Held, S.D.E. & Špinka, M. (2011) Animal play and animal welfare. Animal Behaviour 81, 891–899.

Hobaiter, C. & Byrne, R.W. (2014) The Meanings of Chimpanzee Gestures. Current Biology 24, 1596–1600. Elsevier.

Holler, J. (2022) Visual bodily signals as core devices for coordinating minds in interaction.

Iki, S. & Hasegawa, T. (2020) Face-to-face opening phase in Japanese macaques’ social play enhances and sustains participants’ engagement in subsequent play interaction. Animal Cognition 23, 149–158.

Iki, S. & Hasegawa, T. (2021) Face-to-face configuration in Japanese macaques functions as a platform to establish mutual engagement in social play. Animal Cognition 24, 1179–1189.

Iki, S. & Kutsukake, N. (2022) Play face in Japanese macaques reflects the sender’s play motivation. Animal Cognition.

Kaplan, G. (2024) The evolution of social play in songbirds, parrots and cockatoos -emotional or highly complex cognitive behaviour or both? Neuroscience & Biobehavioral Reviews, 105621.

Kipper, S. & Todt, D. (2002) The use of vocal signals in the social play of barbary macaques. Primates 43, 3–17.

Kuczaj, S.A. & Horback, K.M. (2013) Play and Emotion. In Emotions of Animals and Humans: Comparative Perspectives (eds S. Watanabe & S. Kuczaj), pp. 87–112. Springer Japan, Tokyo.

Lazaro-Perea, C. (2001) Intergroup interactions in wild common marmosets, *Callithrix jacchus*: territorial defence and assessment of neighbours. Animal Behaviour 62, 11–21.

Luo, Y., Stephens, D.A., Verma, A. & Buckeridge, D.L. (2021) Bayesian latent multi-state modeling for nonequidistant longitudinal electronic health records. Biometrics 77, 78–90.

Maglieri, V., Zanoli, A., Mastrandrea, F. & Palagi, E. (2023) Don’t stop me now, I’m having such a good time! Czechoslovakian wolfdogs renovate the motivation to play with a bow. Current Zoology 69, 50–58.

Mancini, G., Ferrari, P.F. & Palagi, E. (2013) In Play We Trust. Rapid Facial Mimicry Predicts the Duration of Playful Interactions in Geladas. PLos One 8, e66481.

Martin, P. & Caro, T.M. (1985) On the Functions of Play and Its Role in Behavioral Development. In Advances in the Study of Behavior (eds J.S. Rosenblatt, C. Beer, M.-C. Busnel & P.J.B. Slater), pp. 59–103. Academic Press.

Mendl, M. (2010) An integrative and functional framework for the study of animal emotion and mood.

Mielke, A. & Carvalho, S. (2022) Chimpanzee play sequences are structured hierarchically as games. Peerj 10, e14294.

Miss, F.M., Adriaense, J.E.C. & Burkart, J.M. (2022) Towards integrating joint action research: Developmental and evolutionary perspectives on co-representation. Neuroscience & Biobehavioral Reviews 143, 104924.

Miss, F.M. & Burkart, J.M. (2018) Corepresentation During Joint Action in Marmoset Monkeys (*Callithrix jacchus*). Psychological Science 29, 984–995.

Mustoe, A.C., Taylor, J.H., Birnie, A.K., Huffman, M.C. & French, J.A. (2014) Gestational cortisol and social play shape development of marmosets’ Hpa functioning and behavioral responses to stressors. Developmental Psychobiology 56, 1229–1243.

Osvath, M. & Sima, M. (2014) Sub-adult Ravens Synchronize their Play: A Case of Emotional Contagion? Animal Behavior and Cognition 2.

Palagi, E. (2008) Sharing the motivation to play: the use of signals in adult bonobos. Animal Behaviour 75, 887–896.

Palagi, E. (2018) Not just for fun! Social playas a springboard for adult social competence in human and non-humanprimates. Behavioral Ecology and Sociobiology 72, 90.

Palagi, E., Burghardt, G.M., Smuts, B., Cordoni, G., DALL’olio, S., Fouts, H.N., Řeháková-Petrů, M., Siviy, S.M. & Pellis, S.M. (2016) Rough-and-tumble play as a window on animal communication. Biological Reviews 91, 311–327.

Pellis, S.M. & Pellis, V.C. (1996) On knowing it’s only play: The role of play signals in play fighting. Aggression and Violent Behavior 1, 249–268.

Pellis, S.M., Pellis, V.C., Ham, J.R. & Stark, R.A. (2023) Play fighting and the development of the social brain: The rat’s tale. Neuroscience & Biobehavioral Reviews 145, 105037.

Petrů, M., Špinka, M., Charvátová, V. & Lhota, S. (2009) Revisiting play elements and self-handicapping in play: A comparative ethogram of five Old World monkey species. Journal of Comparative Psychology 123, 250–263.

Phaniraj, N., Brügger, R.K. & Burkart, J.M. (2024) Marmosets mutually compensate for differences in rhythms when coordinating vigilance. Plos Computational Biology 20, e1012104. Public Library of Science.

R Core Team (2015) R: A language and environment for statistical computing. R Foundation for Statistical Computing. Computing.

Spinka, M., Newberry, R.C. & Bekoff, M. (2001) Mammalian Play: Training for the Unexpected. The Quarterly Review of Biology 76, 141–168. The University of Chicago Press.

Špinka, M., Palečková, M. & Řeháková, M. (2016a) Metacommunication in social play: the meaning of aggression-like elements is modified by play face in Hanuman langurs (Semnopithecus entellus). Behaviour 153, 795–818. Brill.

Špinka, M., Palečková, M. & Řeháková, M. (2016b) Metacommunication in social play: the meaning of aggression-like elements is modified by play face in Hanuman langurs (Semnopithecus entellus). Behaviour 153, 795–818. Brill.

Stan Development Team (2020) RStan: The R interface to Stan. http://mc-stan.org/.

Stevenson, M.F. & Poole, T.B. (1976) An ethogram of the common marmoset (Calithrix jacchus jacchus): General behavioural repertoire. Animal Behaviour 24, 428–451.

Stevenson, M.F. & Poole, T.B. (1982) Playful interactions in family groups of the common marmoset (Callithrix jacchus jacchus). Animal Behaviour 30, 886–900.

Takahashi, D.Y., Narayanan, D.Z. & Ghazanfar, A.A. (2013) Coupled Oscillator Dynamics of Vocal Turn-Taking in Monkeys. Current Biology 23, 2162–2168. Elsevier.

Townsend, S.W., Koski, S.E., Byrne, R.W., Slocombe, K.E., Bickel, B., Boeckle, M., Braga Goncalves, I., Burkart, J.M., Flower, T., Gaunet, F., Glock, H.J., Gruber, T., Jansen, D.A.W.A.M., Liebal, K., Linke, A., Et Al. (2017) Exorcising Grice’s ghost: an empirical approach to studying intentional communication in animals. Biological Reviews 92, 1427–1433.

Voland, E. (1977) Social play behavior of the common marmoset (Callithrix jacchus Erxl., 1777) in captivity. Primates 18, 883–901.

Weigel, E.A. & Berman, C.M. (2018) Body signals used during social play in captive immature western lowland gorillas. Primates 59, 253–265.

Wiltshire, C., Lewis-Cheetham, J., Komedová, V., Matsuzawa, T., Graham, K.E. & Hobaiter, C. (2023) DeepWild: Application of the pose estimation tool DeepLabCut for behaviour tracking in wild chimpanzees and bonobos. Journal of Animal Ecology 92, 1560–1574.

Wright, K.R., Mayhew, J.A., Sheeran, L.K., Funkhouser, J.A., Wagner, Ronald. S., Sun, L.-X. & Li, J.-H. (2018) Playing it cool: Characterizing social play, bout termination, and candidate play signals of juvenile and infant Tibetan macaques (Macaca thibetana). Zoological Research 39, 272–283.

Yanagi, A. & Berman, C.M. (2014a) Body signals during social play in free-ranging rhesus macaques (Macaca mulatta): A systematic analysis. American Journal of Primatology 76, 168–179.

Yanagi, A. & Berman, C.M. (2014b) Functions of multiple play signals in free-ranging juvenile rhesus macaques (Macaca mulatta). Behaviour 151, 1983–2014. Brill.

Yanagi, A. & Berman, C.M. (2017) Does behavioral flexibility contribute to successful play among juvenile rhesus macaques? Behavioral Ecology and Sociobiology 71, 156.

