## Supplemental Info for "Common marmosets use body posture as multi-functional signal to solicit, maintain, and modify social play"

**S1: Group composition**

| **Group** | **Individuals** | **Sex** | **Status** | **Birth date** | **Data collection** |
|---|---|---|---|---|---|
| Washington | Washington | F | breeder | Aug 30, 2013 | Oct – Feb 2021/22 |
| | Lotus | M | breeder | July 3, 2012 | |
| | Wall-e | F | infant | August 12, 2021 | |
| | Woody | M | infant | August 12, 2021 | |
| Jambi | Jambi | F | breeder | Nov 10, 2015 | Sept – Jan 2021/22 |
| | Werewolf | M | breeder | May 10, 2018 | |
| | Jafar | M | infant | July 7, 2021 | |
| | Jaggar | M | infant | July 7, 2021 | |
| Guapa | Guapa | F | breeder | May 4, 2018 | May – Sep 2021 |
| | Ninja | M | breeder | April 12, 2015 | |
| | Gimli | M | infant | March 5, 2021 | |
| | Guarana | M | infant | March 5, 2021 | |

**S2: Data collection**

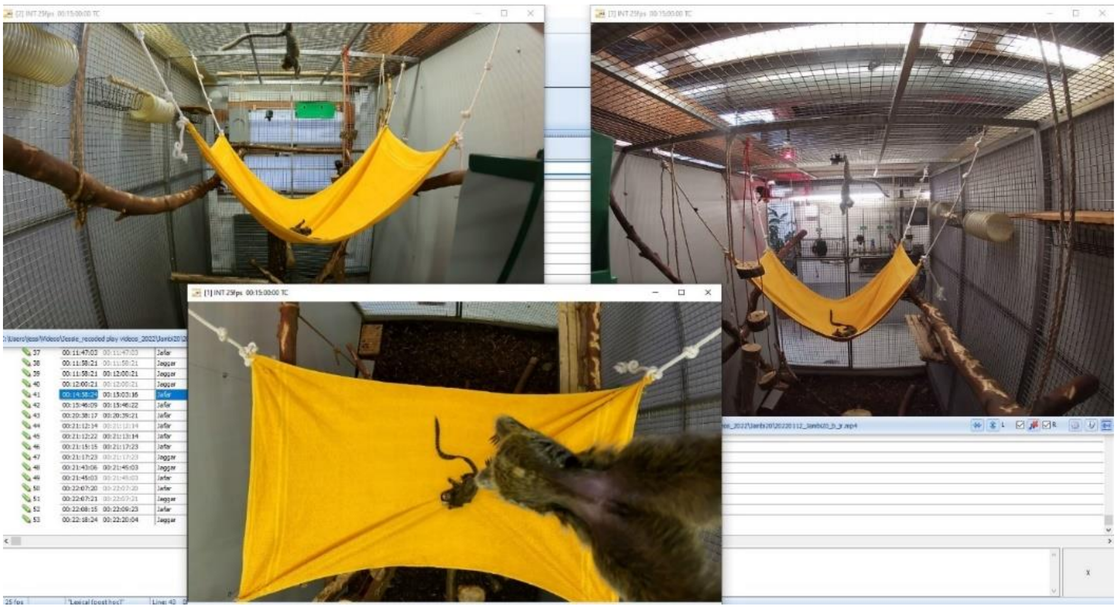

Data collection included a set-up of 3 cameras with a front, back, and top angle

### S3: Ethogram

| | | Definition | Behavioral components | Difference with other types |
|---|---|---|---|---|
| *Play signal* | **Hide** | Focal tries to hide from another individual. The focal does not need to stay hidden in one exact location (e.g. may move along a branch while remaining hidden) or be successful at hiding (i.e. the other may detect where the focal is). During hiding the focal visually fixates on the other and thus often shows their face with the rest of the body hidden. | Hiding | NA |
| | **Stalk** | Focal makes side-ways, jerky, crab-like movements, while fixating visually on another. The focal's tufts are often partially or fully flattened, all feet and hands are flat against the surface, and the focal's body may have a slight dip down of the upper body. | Moving side-ways in a jerky manner | NA |
| | **Supine** | Focal lies on its back or side with full or partial exposure of abdominal/genital area and min. 2 limbs are lifted off the surface. Focal is visually fixated on another individual. In other animals it is also referred to as a self-handicapping posture. | Showing (part of) ventral area | NA |
| *Play type* | **Pounce** | Focal jumps and lands fully or partially on the body of another individual. | Jumping | NA |
| | **Wrestle** | Two individuals are in a full or partial embrace with multiple behavioral components rapidly occurring in sequence. Wrestle occurs without fully biting the other or pinning the other down for a longer time. The coding continues as long as animals are in physical contact (e.g. individuals may look away while performing any of the behaviors). Focal is the individual who establishes first physical contact. | Play biting, grabbing, kicking, pulling, gripping, hitting, pushing, rolling, embracing, or scratching | Different from other play types: if focal does e.g. a pull, and then starts wrestle, code pull first and then wrestle; if pull occurs during wrestle, with no separation of contact, then only code wrestle. |
| | **Pull** | Focal pulls on another individual, by grabbing either the tufts, tail, or limbs. Pulling is defined as exerting force to cause movement towards oneself. | Pulling | Different from touch: pull has an exerting force versus touch having a stagnant focus |
| | **Chase** | Focal runs behind another subject, who runs away from the focal, and both run in parallel. The chase is in one direction only, with the focal of the chase in the back (the chaser), and the receiver of the chase in the front (chasee). When the chase turns around, and thus in the reverse direction, the focal becomes receiver, the receiver becomes focal, and a new chase is coded. | Running, leaping | Different from catch: chase is uni-directional and catch is bi-directional. Chase may also include multiple locations and catch remains in one location. |
| | **Touch** | Focal touches another subject once with one of the forelimbs. | Slapping, batting, hitting, or placing one of hand on the other | Different from wrestle: when touch leads immediately to wrestle, then there is no separate code for touch but only for wrestle |
| | **Catch** | Focal and other individual make short back-and-forth movements toward each other, visually fixating one another, remaining in one location (i.e. bi-directional movement), and using forelimbs to slap or grab the other. Looks like the subjects are pretending to grab the other but not fully succeeding. | Bi-directional partial grabbing | Different from wrestle: body contact during catch is only through slapping or pretend-grabbing of the other; during wrestle there is always body contact |
