## Supplemental Stats for "Common marmosets use body posture as multi-functional signal to solicit, maintain, and modify social play"

### Statistical modelling supplement for: Common marmosets use body posture as multi-functional signal to solicit, maintain, and modify social play

**2 Model of duration of play ~ signal (Prediction 2) 7**

**3 Model of diversity of play ~ signal (Prediction 3) 10**

**4 Model of intensity of play ~ signal (Prediction 4) 10**

**5 Model of turn-taking in sending/receiving (Exploratory question 1) 11**

**6 Model of the number of dyads (Exploratory question 2) 12**

**References 12**

### 1 Multi-state time-to-event model (Prediction 1)

#### 1.1 Motivation

Multi-state time-to-event models (also called survival models) are a popular method for analyzing both the duration of behavioral states and the state-dependent transitions between behaviors (Andersen, Abildstrom, and Rosthøj 2002; Meira-Machado et al. 2009; Gygax, Zeeland, and Rufener 2022; Le-Rademacher, Therneau, and Ou 2022). Imagine that an animal can be in only three states: foraging, grooming, or nesting. One might ask (1) how long will they spend in the nesting state, and (2) when they stop nesting, will their next behavior be foraging or grooming? Typically, such models assume a first-order Markov process: the duration and type of transition depends only on the current state, not on the past states. For example, the time spent foraging is independent of how much time the animal had previously spent nesting. Similarly, if the animal takes a break from nesting to groom themselves, then the memory of nesting is "forgotten" and cannot influence subsequent behavior–all that matters is that they are grooming now.

There are many ways that the Markov assumption can be violated, motivating more complex models and higher-degree Markov models that have longer memories (Langeheine and Van de Pol 2000; Ching, Fung, and Ng 2004). But these models can be challenging to fit and interpret due to data sparsity. A general challenge in the application of these models to behavioral data is how to treat the space *between* behavioral states. After the animal has stopped grooming itself–but before it starts foraging–there is a lag or transitional period, in which their behavior is neither state. It is tempting to classify this as a state in an of itself, perhaps labelled "inactive" or "rest". But this naive approach breaks the dependency between grooming and foraging. From the model's point-of-view, the time until foraging starts is now independent of grooming, artificially disrupting the behavioral sequence. Conversely, if a lag between behaviors is sufficiently long, then it may be a "true" break in the sequence, in which case the future behavior should be treated as independent of the past. In the context of our study of marmoset behavior, we have two observable behavioral states (signal and play) and one latent state (rest).

In practice, researchers may use heuristics to define behavioral "bouts" which are assumed to be independent of each other if a long enough time has elapsed (e.g., 10 seconds). However, this use of thresholds is quite subjective, and if the distribution of lags is not clearly bimodal, this procedure introduces measurement error as some lags are incorrectly classified as resting states. To address this problem, we adopt a mixture model approach that allows us to probabilistically classify each lag as either an brief pause (in which case the behavior sequence is uninterrupted) or a long rest (in which case a rest state interrupts the sequence). In this sense, our model is a special case of a Hidden Markov Model: rest states are hidden but non-resting states are directly observable. In the next section, we outline our assumed generative model for both the latent states and observable data.

#### 1.2 Generative Model

In this section we describe a (partially) generative continuous-time Markov model of marmoset behavioral states. It is only partially generative because we omit: (1) the initial states of the behavioral sequences and (2) the lags between behaviors of the same state (e.g., two different types of play punctuated by a lag, but each belonging to the macro-state of "play"). The latter depends on many factors such as how the ethogram is constructed (i.e., how to categorize the diversity of behavioral phenotypes) and the decisions of the behavioral coder.

##### 1.2.1 Latent states

For each behavioral state at time $t > 1$, the duration of some state $y \in [1, K]$ (i.e., the waiting time) is denoted $x_t$.

$$
\begin{aligned}
&x_t \sim \text{Exponential}(\sum_{k=1}^{K} \lambda_{[y_t,\ k]}\ ) \\
&y_{t+1} \mid x_t \sim \text{Categorical}(\boldsymbol{p}_t) \\
&p_{tk} \propto \lambda_{[y_{t-1},k]} S(x_t) \\
&S(x_t) = \prod_{j=1}^{K} e^{-\lambda_{[y_{t-1},j]} x_t} = e^{-\sum_{k=1}^{K} \lambda_{[y_{t-1},\ k]} x_t}
\end{aligned}
\tag{1}
$$

Where $\lambda_{[i,j]}$ denotes the rate of transitions from state $i$ to state $j$ that fills the off-diagonals of a transition matrix $\boldsymbol{\Lambda}$. The diagonal elements of $\boldsymbol{\Lambda}$ is fixed to 0. After sampling the duration, we can sample the new behavioral state $y_{t+1}$. The probability of transitioning to state $k$ is proportional to ($\propto$) the probability that the transition from $y_t$ to $k$ happened after duration $x_t$ AND the probability that no other transitions occurred before $x_t$, the latter represented by the survival function $S(x_t)$. Also note that $p_{t[k\ =\ t-1]} = 0$ because some transition must have happened by time $x_t$.

The last component of our generative model is the duration of the pause between consecutive non-rest states. $\zeta$ ("zeta") can be interpreted as the "lag" between two behaviors, in which the subject is doing neither $y_t$ nor $y_{t+1}$–but is also not in a true rest state.

$$
\zeta_{t+1} = \begin{cases} 0, & \text{if } k = 1 = \text{rest} \\ \text{Exponential}(\lambda_\zeta), & \text{otherwise} \end{cases}
\tag{2}
$$

Thus, for each time step $t > 1$ we generate a triplet $(x_t, y_{t+1}, \zeta_{t+1})$, where $\zeta_{t+1} = 0$ if $y_t$ was a rest state. The initial state $y_{t=1}$ might be sampled from the stationary distribution, or modeled separately. To generate samples from this model, one also needs to assign some prior distributions to the parameters. Additionally, to ensure that the resting states are longer on average than the pause states, we impose an ordered constraint on the parameters such that $\sum_{k=2}^{K} \lambda_{[1,\ k]} < \lambda_{\zeta}$. In practice, we use a Dirichlet decomposition to enforce this constraint: $\lambda_{[1,k]} = \sigma(\boldsymbol{z})_k \phi \lambda_{\zeta}$. Where $\sigma()$ denotes the softmax function evaluating a vector of latent scores for $K-1$ state transitions, $\phi \in (0,1)$ controls the total rate of transitions out of the rest state, and $\lambda_{\zeta}$ is the rate of the Exponential distribution for pauses. As $\phi \to 1$, the transition rate from the rest state approaches the speed of pauses. By bounding this parameter, we ensure that the mixture components are identified such that $\lambda_{\zeta}$ is strictly greater than the total rate of transitions out of rest.

##### 1.2.2 Observational model

In our study we observe $\{x_{t-1}, y_{t-1}, y_t, \zeta_t\}$ for each time $t > 1$. Furthermore, $y_{t-1}, y_t$ are always $k > 1$ because rest states can only be inferred rather directly observed. We formalize this process in the algorithm below:

**Algorithm 1:** Data Generating Process for Multi-state Model

**Data:** Initial state $y_1$, parameters $\{\boldsymbol{\Lambda},\ \lambda_{\zeta}\}$, target duration $\mathrm{T}_{\max}$

**Result:** Data $\{(x_t, y_t, \zeta_t)\}_{t=1}^{T}$ where $T$ is the number of discrete time steps recorded until total duration $\mathrm{T}_{\max}$ is reached

**Function** `GenerateData`($y_1, \boldsymbol{\Lambda}, \lambda_{\zeta}, T_{max}$)**:**

- $t \leftarrow 1$ ;
- **while** $\sum_{i=1}^{t}(x_{i-1} + \zeta_i) < T_{max}$ **do**
  - Generate $x_t \sim \text{Exponential}\left(\sum_{k=1,\ k \neq y_t}^{K} \lambda_{[y_t,\ k]}\right)$;
  - Generate $y_{t+1} \mid x_t \sim \text{Categorical}(\boldsymbol{p}_t)$;
  - **if** $y_{t+1} > 1$ **then**
    - Generate $\zeta_{t+1} \sim \text{Exponential}(\lambda_{\zeta})$;
  - **else**
    - Generate $x_{\text{temp}} \sim \text{Exponential}\left(\sum_{k=1,\ k \neq y_t}^{K} \lambda_{[y_t,\ k]}\right)$;
    - Generate $y_{t+1} \mid x_{\text{temp}} \sim \text{Categorical}(\boldsymbol{p}_t)$, replacing the previously generated value;
    - Update $\zeta_{t+1} \leftarrow x_{\text{temp}}$;
  - Record $(x_t, y_{t+1}, \zeta_{t+1})$;
  - Update $t \leftarrow t + 1$;
- **return** Data $\{(x_t, y_t, \zeta_t)\}_{t=1}^{T}$

##### 1.3 Likelihood

Each $\zeta$ could have been generated by either a transition from a rest state ($k = 1$) or by a pause between non-rest states ($k > 1$). In other words, at each time point the data could have been generated by either one transition ($y_t \to y_{t+1}$) or two transitions ($y_t \to \text{rest} \to y_{t+1}$). Therefore the likelihood is defined as a finite mixture model:

$$p(x_{t-1}, y_t, \zeta_t \mid y_{t-1}, \mathbf{\Lambda}, \lambda_\zeta) = \begin{cases} \lambda_{y_{t-1},1} S(x_{t-1} \mid \mathbf{\Lambda}) \times \lambda_{1,y_t} S(\zeta_t \mid \mathbf{\Lambda}) + \\ S(x_{t-1} \mid \mathbf{\Lambda}) \times \lambda_\zeta S(\zeta_t \mid \lambda_\zeta) & \text{if } y_t = y_{t-1}, \text{ and} \\ \\ \lambda_{y_{t-1},1} S(x_{t-1} \mid \mathbf{\Lambda}) \times \lambda_{1,y_t} S(\zeta_t \mid \mathbf{\Lambda}) + \\ \lambda_{y_{t-1},y_t} S(x_{t-1} \mid \mathbf{\Lambda}) \times \lambda_\zeta S(\zeta_t \mid \lambda_\zeta) & \text{if } y_t \neq y_{t-1}. \end{cases} \tag{3}$$

For each case above, the first line denotes the the likelihood of the data given that the lag duration $\zeta_t$ was generated by an intermediate rest state. The second line denotes the likelihood given that $\zeta_t$ was generated by a brief pause between $y_{t-1}$ and $y_t$. The only difference between the two cases ($y_t = y_{t-1}$, $y_t \neq y_{t-1}$) is that in the former, the time at transition is right-censored.

Due to the memoryless property of the exponential distribution, we do not need to keep track of repeated self-transitions (e.g., $y_t = y_{t-1} = y_{t-2}$). For example, imagine a sequence of two play behaviors with duration $x_a$ and $x_b$, respectively, broken up only by a pause. In this case the appropriate survival function is $S(x_a + x_b)$, reflecting the total duration of the play state. But given the uncertainty around rest states, doing so complicates the likelihood as we need to marginalize over $2^m$ possible explanations of the data (where $m$ is the number of sequential behaviors of the same state). Conveniently, the instantaneous rate of transition in the Exponential distribution is constant over time such that $S(x_a + x_b) = S(x_a) * S(x_b)$, as can also be seen in Eq 1. This justifies the simplified likelihood defined above, but it is important to note that it would not generalize to other distributions, such as the Gamma or Weibull.

##### 1.4 Classification as Pause or Rest

We can calculate the posterior probability that a transition was generated by a rest state using Bayes' rule:

$$p(\text{rest}|x_{t-1}, y_t, \zeta_t, y_{t-1}, \mathbf{\Lambda}, \lambda_\zeta) = \begin{cases} \frac{\lambda_{y_{t-1},1} S(x_{t-1}|\mathbf{\Lambda}) \times \lambda_{1,y_t} S(\zeta_t|\mathbf{\Lambda})}{\lambda_{y_{t-1},1} S(x_{t-1}|\mathbf{\Lambda}) \times \lambda_{1,y_t} S(\zeta_t|\mathbf{\Lambda}) \;+\; S(x_{t-1}|\mathbf{\Lambda}) \times \lambda_\zeta S(\zeta_t|\lambda_\zeta)} & \text{if } y_t = y_{t-1}, \text{ and} \\ \\ \frac{\lambda_{y_{t-1},1} S(x_{t-1}|\mathbf{\Lambda}) \times \lambda_{1,y_t} S(\zeta_t|\mathbf{\Lambda})}{\lambda_{y_{t-1},1} S(x_{t-1}|\mathbf{\Lambda}) \times \lambda_{1,y_t} S(\zeta_t|\mathbf{\Lambda}) \;+\; \lambda_{y_{t-1},y_t} S(x_{t-1}|\mathbf{\Lambda}) \times \lambda_\zeta S(\zeta_t|\lambda_\zeta)} & \text{if } y_t \neq y_{t-1}. \end{cases} \tag{4}$$

Because there are only two mixture components, $p(\text{pause}) = 1 - p(\text{rest})$.

#### 1.5 Multilevel Model

We extend the basic model to accommodate repeated measures at the session level. For each session $s$: $\lambda_{s_{[i,j]}} = \exp(\log(\lambda_{[i,j]}) + \nu_{[s,i,j]})$. Where $\boldsymbol{\nu}$ is an array of random effects that vary at the group level with common standard deviation hyper-parameters $\boldsymbol{\sigma}_{\text{session}}$ and session-level correlations between parameters $\boldsymbol{\Omega}_{\text{session}}$. We also allow the rate of pauses $\lambda_\zeta$ to vary depending on the current behavioral state, and introduce a hurdle component (i.e., "zero inflation") to the lag model to account for durations of length 0, which can occur in the case of temporally overlapping behaviors.

##### 1.5.1 Priors

As a pre-processing step, the duration of behavior and lags were scaled by the average behavioral duration. This was done to facilitate model convergence by keeping the parameters near the unit scale, which aids in the convergence of MCMC chains (Team 2024).

$$
\begin{aligned}
\lambda &\sim \mathcal{N}(0,2) && \text{non-rest rates} \\
\lambda_\zeta &\sim \mathcal{N}(0,2) && \text{pause rates} \\
z &\sim \mathcal{N}(0,1) && \text{latent-scale rest transitions} \\
\phi &\sim \text{Beta}(1,1) && \text{proportional rate of rest, relative to pause} \\
\theta_{\text{pause}} &\sim \text{Beta}(1,1) && \text{probability of zero-duration lag}
\end{aligned}
$$

$$
\begin{aligned}
\nu &\sim \mathcal{N}(0,\boldsymbol{\Sigma}) && \text{session-level random effects} \\
\boldsymbol{\Sigma} &= \text{diag}(\boldsymbol{\sigma}_{\text{session}})\,\boldsymbol{\Omega}_{\text{session}}\,\text{diag}(\boldsymbol{\sigma}_{\text{session}}) \\
\sigma_{\text{session}} &\sim \text{Exponential}(2) \\
\boldsymbol{\Omega}_{\text{session}} &\sim \text{LKJ}(2)
\end{aligned}
$$

#### 1.6 Fitting the Model

We implemented the multistate model described above to our study data using the Stan probabilistic programming language (Carpenter et al. 2017) via the cmdstanr R package (Gabry et al. 2024), which fits Bayesian models using Hamiltonian Markov Chain Monte Carlo. We ran 4 parallel chains for 500 iterations of warmup and 5000 iterations of sampling. Markov chain convergence was assessed using standard diagnostics (no divergent transitions, the number of effective samples, the Gelman-Rubin diagnostic, and visual inspection of trace plots). See Figure 1 for trace plots of key model parameters. Stan code for all models is included in the supplementary materials. `msm_classifier_zi.stan` is the file for the multistate model described in this section.

##### 1.7 Model Checking

Model checking is subtle in this context, because the total duration of behavioral states is inferred from the data itself. However, it is possible to check the predicted duration of lags between states compared to the observed duration. Figure 2 displays posterior predictive checks for the distribution of lags at the session level.

##### 1.8 Using the Classifier to Sample Behavioral Sequences

Equation 4 gives us the probability that each lag was generated by either a rest or a pause. These probabilities jointly imply a distribution of behavioral sequences, reflecting uncertainty in the classifications. To appropriately propagate this uncertainty into subsequent analyses, we used drew 100 samples. For all of the subsequent analyses that rely on these classifications, we fit the models 100 times (to each sampled set of behavioral sequences) and combined the resulting samples. The next sections provide model definitions for these additional analyses.

#### 2 Model of duration of play ∼ signal (Prediction 2)

For each play duration $i$ in session $s$ preceded by a signal (signal = 1) or not (signal = 0):

$$
\begin{aligned}
&\text{duration}_{i,s} \sim \text{Gamma}(\alpha, \frac{\alpha}{\mu_{i,s}}) \\
&\mu_{i,s} = \exp(\beta_0 + \nu_{0_s} + (\beta_{\text{signal}} + \nu_{\beta_s})\text{signal}_i) \\
&\alpha \sim \text{HalfNormal}(1, 2) \\
&\beta_0, \beta \sim \mathcal{N}(0, 2) \\
&\nu_{0_s} \sim \mathcal{N}(0, \sigma_{\text{session}_0}) \\
&\nu_{\beta_s} \sim \mathcal{N}(0, \sigma_{\text{session}_\beta}) \\
&\sigma_{\text{session}_0}, \sigma_{\text{session}_\beta} \sim \text{Exponential}(1)
\end{aligned}
$$

To contrast the two conditions (rest before vs signal before), we take the difference in $\mu$ for each condition, marginalizing over session-level variation.

The Stan file for this model is `duration_model_gamma.stan`.

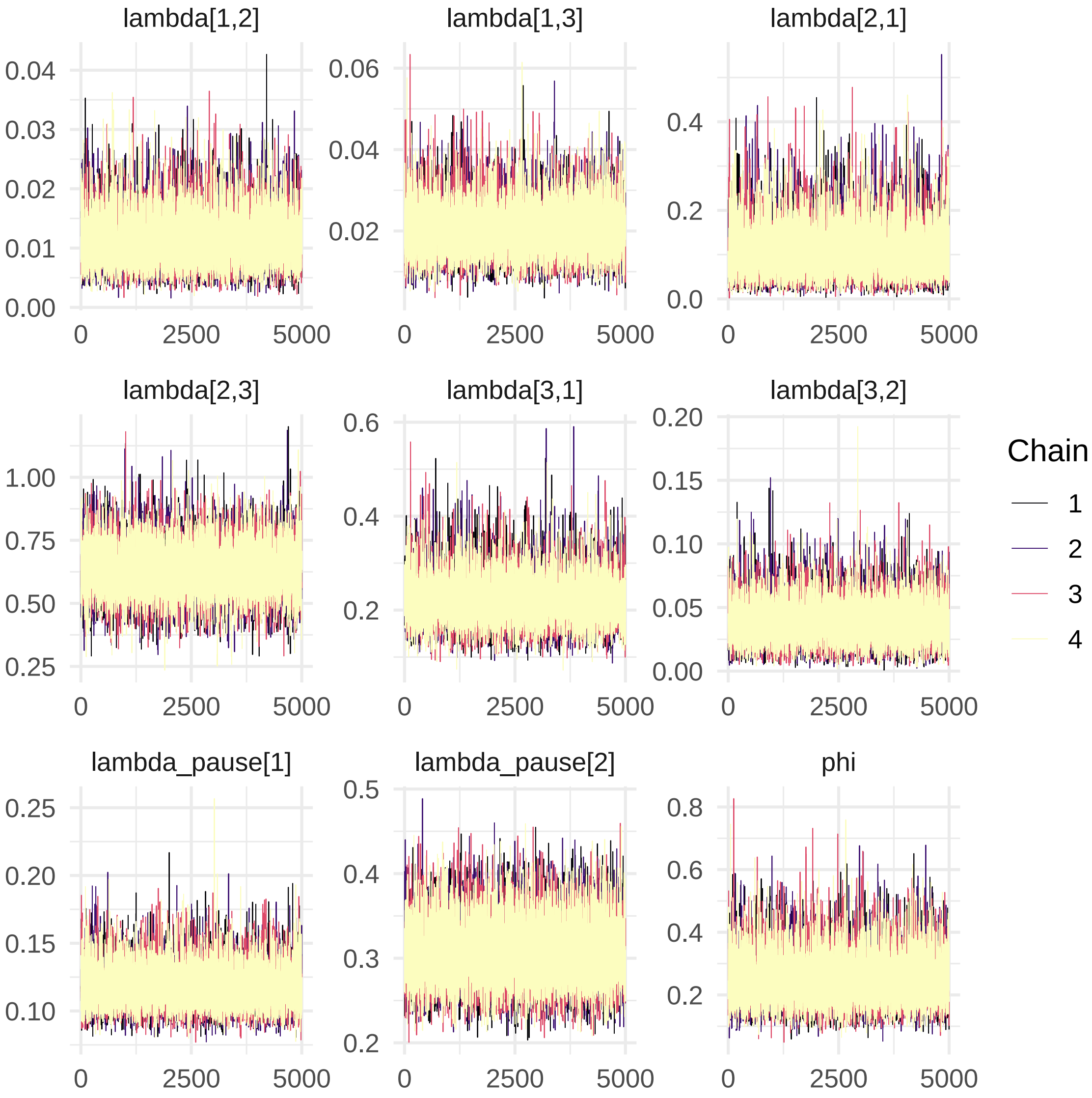

Figure 1: Traceplots demonstrating convergence and mixing of post-warmup samples.

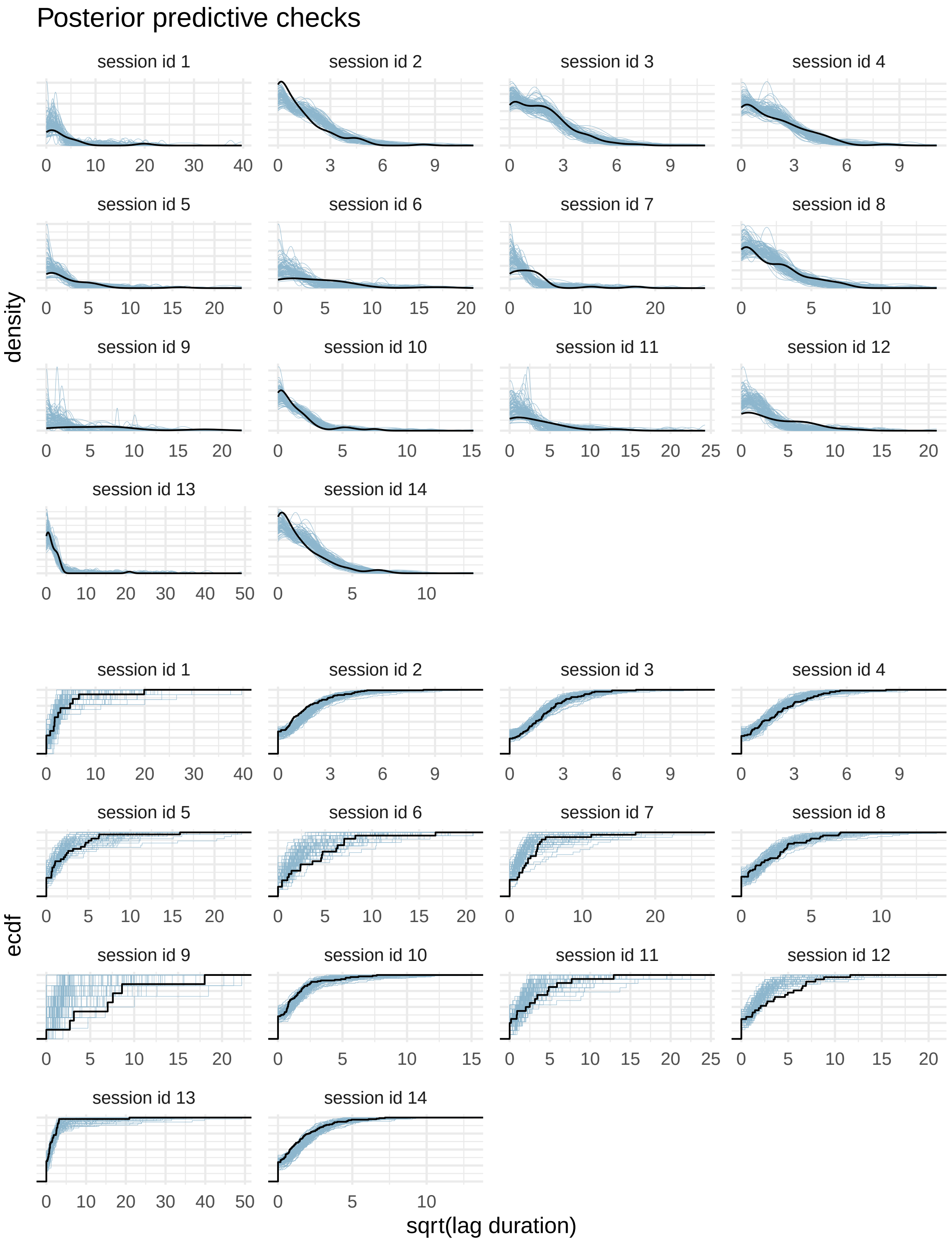

Figure 2: Posterior predictive checks, grouped by session. Shown on the square root scale for enhanced visualization.

#### 3 Model of diversity of play ~ signal (Prediction 3)

For each count of unique play behaviors (diversity) within play state $i$ in session $s$ preceded by a signal (signal = 1) or not (signal = 0):

$$
\begin{aligned}
&\text{diversity}_{i,s} \sim \text{OrderedLogistic}(\eta_{i,s}, \boldsymbol{c_i}) \\
&\eta_{i,s} = \nu_{0_s} + (\beta + \nu_{\beta_s})\text{signal}_i \\
&\boldsymbol{c_i} = \begin{cases} \boldsymbol{c}_{\text{rest}} & \text{if signal}_i = 0, \text{ and} \\ \boldsymbol{c}_{\text{signal}} & \text{if signal}_i = 1. \end{cases} \\
&\beta \sim \mathcal{N}(0, 2) \\
&\boldsymbol{c}_{\text{rest}}, \boldsymbol{c}_{\text{rest}} \sim \mathcal{N}(0, 2) \\
&\nu_{0_s} \sim \mathcal{N}(0, \sigma_{\text{session}_0}) \\
&\nu_{\beta_s} \sim \mathcal{N}(0, \sigma_{\text{session}_\beta}) \\
&\sigma_{\text{session}_0}, \sigma_{\text{session}_\beta} \sim \text{Exponential}(1)
\end{aligned}
$$

Estimating a different set of cutpoints $\boldsymbol{c}$ for both the "signal before" and "rest before" conditions relaxes the parameteric assumptions of the ordinal model implied by using a common set of cutpoints. To contrast the two conditions, we calculated the expected value (on the original measurement scale) for each condition and take the difference, marginalizing over session-level variation.

The additional model of play diversity that adjusts for duration is nearly the same as the defined above, simply adding the standardized duration of the play state as a linear predictor with prior $\beta_{\text{duration}} \sim \mathcal{N}(0, 2)$. The Stan files for these models are `diversity_model_ord.stan` and `diversity_model_ord_duration.stan`.

#### 4 Model of intensity of play ~ signal (Prediction 4)

We model the presence or absence of each behavior type $j$ within play state $i$ in session $s$:

$$
\begin{aligned}
&\text{behavior}_{j,i,s} \sim \text{Bernoulli}(p_{j,i,s}) \\
&p_{j,i,s} = \text{logit}^{-1}(\beta_{0_j} + \nu_{0_{j,s}} + (\beta_{\text{signal}_j} + \nu_{\beta_{j,s}})\text{signal}_i) \\
&\beta_{0_j}, \beta_j \sim \mathcal{N}(0, 2) \\
&\nu_{0_{j,s}} \sim \mathcal{N}(0, \sigma_{\text{session}_{0_j}}) \\
&\nu_{\beta_{j,s}} \sim \mathcal{N}(0, \sigma_{\text{session}_{\beta_j}}) \\
&\sigma_{\text{session}_{0_j}}, \sigma_{\text{session}_{\beta_j}} \sim \text{Exponential}(1)
\end{aligned}
$$

From this model, we calculate behavior-specific contrasts between the "rest before" (signal = 0) and "signal before" (signal = 1) conditions, including those for wrestle and chase. We compute the difference in the probability of the behavior occurring between conditions, marginalizing over session-level variation.

The Stan file for this model is `behavior_model.stan`.

### 5 Model of turn-taking in sending/receiving (Exploratory question 1)

For each observed signal $\rightarrow$ play sequence in session $s$, we categorize the roles taken by the constituent dyads as:

role = 1: roles retained (the sender of the signal is the initiator of play)

role = 2: roles reversed (the receiver of the signal is the initiator of play)

role = 3: other (the sender and/or receiver changes identity between the signal and play state)

Which gives $K = 3$ response categories. The probability of each category is denoted $\theta_k$.

$$
\begin{aligned}
&\text{role}_s \sim \text{Categorical}(\boldsymbol{\theta}_s) \\
&\boldsymbol{\theta}_s = \sigma(z_s) \\
&z_{1,s} = \beta_{0_1} + \nu_{0_{1,s}} \\
&z_{2,s} = \beta_{0_2} + \nu_{0_{2,s}} \\
&z_{3,s} = 0 \\
&\beta_{0_k} \sim \mathcal{N}(0, 1) \\
&\nu_{0_{k,s}} \sim \mathcal{N}(0, \sigma_{\text{session}_{0_k}}) \\
&\sigma_{\text{session}_{0_k}} \sim \text{Exponential}(1)
\end{aligned}
$$

Where $\sigma()$ denotes the softmax function. We calculate the difference in the probability of each role type via the difference in $\theta_k$, marginalizing over session-level variation.

The Stan file for this model is `roles_model.stan`.

### 6 Model of the number of dyads (Exploratory question 2)

We model the number of unique dyads (ndyads) that participated in a play state $i$ in session $s$ preceded by a signal (signal = 1) or not (signal = 0):

$$
\begin{aligned}
&\text{ndyads}_{i,s} \sim \text{OrderedLogistic}(\eta_{i,s}, \boldsymbol{c_i}) \\
&\eta_{i,s} = \nu_{0_s} + (\beta + \nu_{\beta_s})\text{signal}_i \\
&\boldsymbol{c_i} = \begin{cases} \boldsymbol{c}_{\text{rest}} & \text{if signal}_i = 0, \text{ and} \\ \boldsymbol{c}_{\text{signal}} & \text{if signal}_i = 1. \end{cases} \\
&\beta \sim \mathcal{N}(0, 2) \\
&\boldsymbol{c}_{\text{rest}}, \boldsymbol{c}_{\text{rest}} \sim \mathcal{N}(0, 2) \\
&\nu_{0_s} \sim \mathcal{N}(0, \sigma_{\text{session}_0}) \\
&\nu_{\beta_s} \sim \mathcal{N}(0, \sigma_{\text{session}_\beta}) \\
&\sigma_{\text{session}_0}, \sigma_{\text{session}_\beta} \sim \text{Exponential}(1)
\end{aligned}
$$

To contrast the two conditions, we calculated the expected value (on the original measurement scale) for each condition and take the difference, marginalizing over session-level variation.

The Stan file for this model is `ndyad_model_ord.stan`.

#### References

Andersen, Per Kragh, Steen Z Abildstrom, and Susanne Rosthøj. 2002. "Competing Risks as a Multi-State Model." *Statistical Methods in Medical Research* 11 (2): 203–15.

Carpenter, Bob, Andrew Gelman, Matthew D Hoffman, Daniel Lee, Ben Goodrich, Michael Betancourt, Marcus A Brubaker, Jiqiang Guo, Peter Li, and Allen Riddell. 2017. "Stan: A Probabilistic Programming Language." *Journal of Statistical Software* 76.

Ching, Wai Ki, Eric S Fung, and Michael K Ng. 2004. "Higher-Order Markov Chain Models for Categorical Data Sequences." *Naval Research Logistics (NRL)* 51 (4): 557–74.

Gabry, Jonah, Rok Češnovar, Andrew Johnson, and Steve Bronder. 2024. "Cmdstanr: R Interface to 'CmdStan'." https://mc-stan.org/cmdstanr/.

Gygax, Lorenz, Yvonne RA Zeeland, and Christina Rufener. 2022. "Fully Flexible Analysis of Behavioural Sequences Based on Parametric Survival Models with Frailties—a Tutorial." *Ethology* 128 (2): 183–96.

Langeheine, Rolf, and Frank Van de Pol. 2000. "Fitting Higher Order Markov Chains." *Methods of Psychological Research Online* 5 (1): 32–55.

Le-Rademacher, Jennifer G, Terry M Therneau, and Fang-Shu Ou. 2022. "The Utility of Multistate Models: A Flexible Framework for Time-to-Event Data." *Current Epidemiology Reports* 9 (3): 183–89.

Meira-Machado, Luís, Jacobo de Uña-Álvarez, Carmen Cadarso-Suárez, and Per K Andersen. 2009. "Multi-State Models for the Analysis of Time-to-Event Data." *Statistical Methods in Medical Research* 18 (2): 195–222.

Team, Stan Development. 2024. *Stan User's Guide* (version 2.35). https://mc-stan.org/docs/stan-users-guide/index.html.
